# The first transchromosomic rat model with human chromosome 21 shows robust Down syndrome features

**DOI:** 10.1101/2021.10.03.462958

**Authors:** Yasuhiro Kazuki, Feng J. Gao, Miho Yamakawa, Masumi Hirabayashi, Kanako Kazuki, Naoyo Kajitani, Sachiko Miyagawa-Tomita, Satoshi Abe, Makoto Sanbo, Hiromasa Hara, Hiroshi Kuniishi, Satoshi Ichisaka, Yoshio Hata, Moeka Koshima, Haruka Takayama, Shoko Takehara, Yuji Nakayama, Masaharu Hiratsuka, Yuich Iida, Satoko Matsukura, Naohiro Noda, Yicong Li, Anna J. Moyer, Bei Cheng, Nandini Singh, Joan T. Richtsmeier, Mitsuo Oshimura, Roger H. Reeves

## Abstract

Progress in earlier detection and symptom management has increased life expectancy and quality of life in people with Down syndrome (DS). However, no drug has been approved to help individuals with DS live independently and fully. Although rat models could support more robust physiological, behavioral, and toxicology analysis than mouse models during preclinical validation, no DS rat model is available due to technical challenges. We developed the first transchromosomic rat model of DS, TcHSA21rat, which contains a freely segregating, EGFP-inserted, human chromosome 21 (HSA21) with >93% of its protein coding genes. RNA-Seq of neonatal forebrains demonstrates that TcHSA21rat not only expresses HSA21 genes but also has an imbalance in global gene expression. Using EGFP as a marker for trisomic cells, flow cytometry analyses of peripheral blood cells from 361 adult TcHSA21rat animals show that 81% of animals retain HSA21 in >80% of cells, the criterion for a “Down syndrome karyotype” in people. TcHSA21rat exhibits learning and memory deficits and shows increased anxiety and hyperactivity. TcHSA21rat recapitulates well-characterized DS brain morphology, including smaller brain volume and reduced cerebellar size. In addition, the rat model shows reduced cerebellar foliation, a prominent feature of DS that is not observed in DS mouse models. Moreover, TcHSA21rat exhibits anomalies in craniofacial morphology, heart development, husbandry, and stature. TcHSA21rat is a robust DS animal model that can facilitate DS basic research and provide a unique tool for preclinical validation to accelerate DS drug development.

## Introduction

Down syndrome (DS), caused by trisomy for human chromosome 21 (HSA21), is the most common viable aneuploidy and occurs in ∼ 1/800 live births^1^. People with DS have varying levels of intellectual disability. They are at high risk of developing other health conditions, such as congenital heart defects (CHD), hearing and vision loss, leukemia, gastrointestinal disease, and early-onset dementia^2, 3^. Significant progress in prenatal detection, symptom management, and awareness of DS has substantially increased life expectancy and quality of life in people with DS^4, 5^. However, most people with DS cannot live independently. Pharmacological approaches to ameliorate different aspects of this complex genetic problem, many tested in mouse models, hold promise, but further research exploring DS mechanisms in optimal model systems is necessary to advance treatments.

HSA21 comprises 46.7 million base pairs (Mb) of DNA (GRCh38.p12). Seventeen and 213 protein coding genes (PCGs) are annotated in the short arm (HSA21p) and long arm (HSA21q), respectively, in addition to 601 annotated non-PCGs. Trisomy changes expression for HSA21 genes at dosage imbalance with the consequence of genome-wide expression imbalance^3, 6^. The first viable DS mouse model, Ts65Dn, contains an extra freely segregating chromosome with 92 of 160 non-keratin associated protein (non-*KRTAP*) genes orthologous to HSA21 and shows features comparable to various DS manifestations, including cognitive impairment, retrusion of the midface skeleton, resistance to solid tumors and cerebellar hypoplasia^7–9^.

Genetic criteria for an optimal DS animal model include: 1) aneuploidy, an extra freely segregating chromosome introducing an extra centromere into every cell; 2) a large fraction of trisomic HSA21 orthologous genes; 3) few or no trisomic genes that are not HSA21 genes/orthologs; 4) no regions of monosomy, as can occur in translocations^10^; and 5) minimal mosaicism. Ts65Dn contains a freely segregating chromosome but is trisomic for a number of non-HSA21 orthologs^10, 11^. Models with direct duplications of conserved regions, such as Dp(16)1Yey, do not have an extra chromosome and centromere^12, 13^. Tc1 was the first transchromosomic aneuploid model and is trisomic for a somewhat rearranged HSA21 carrying ∼75% of PCGs but shows extensive mosaicism^14^. TcMAC21 is currently the most complete genetic mouse model of DS and contains a hybrid chromosome “HSA21q-MAC” composed of a mouse artificial chromosome (MAC) vector engineered with HSA21q^15^. TcMAC21 has little or no mosaicism and carries 93% of HSA21 PCGs and no non-HSA21 human genes.

A study of 59 new FDA-approved drugs from 2015 to 2016 shows that placebo-controlled or drug comparator clinical trials have an average cost of $35.1 million^16^. Drugs for treating central nervous system disorders have low approval rates^17^. The use of trisomic DS mouse models has identified several potential therapeutic candidates to improve learning and memory in people with trisomy 21^18–22^. Animal models that support more robust physiological and behavioral analysis will provide an essential layer of preclinical validation, with improved outcomes for drug development.

The laboratory rat, *Rattus norvegicus*, was the first mammalian species domesticated for scientific research in 1828 and had the third complete mammalian genome sequence in 2004^23^. Humans and rodents separated from a common ancestor ∼ 75 million years (Myr) ago, while mice and rats diverged ∼ 12-24 Myr^23–25^. Rats have 21 pairs of chromosomes, compared with 23 in humans and 20 in mice. The rat genome (Rnor_6.0) comprises ∼2.87 gigabases (Gb), somewhat smaller than the human genome (∼3.1 Gb, GRCh38.p12) and slightly larger than the mouse genome (2.73 Gb, GRCm38.p6). The larger body and organ size of rats permits better imaging and surgical/physiological interventions than in mice. Rat organ morphology such as cerebellar foliation is more similar to the human than is that of the mouse^26^. Rat behaviors are well characterized and nuanced, including more affiliative social behavior, and rats are generally competent in and less stressed by cognitive tests like the Morris water maze (MWM)^27, 28^. Moreover, rats are the predominant model system in safety and toxicology studies of new compounds or nutrients. Here, we developed and characterized the first transchromosomic (Tc) rat model of DS, “TcHSA21rat”, which contains a substantially intact and freely segregating EGFP-inserted HSA21 and recapitulates many phenotypes associated with DS.

## Results

### The generation and whole genome sequencing (WGS) of TcHSA21rat

The hybrid A9 cells containing a copy of HSA21 were generated previously^29^. Because DT40 cells have a high frequency of homologous recombination, the HSA21 was moved from A9 into DT40 cells through microcell-mediated chromosome transfer (MMCT)^30, 31^. In hybrid DT40 cells, the HSA21 was modified with a loxP site at its near centromere gene-free locus, “NC_000021.9” (from 13,021,348 bp to 13,028,858 bp), to form the “HSA21-loxP” chromosome^30^. In this study, the HSA21-loxP chromosome was then transferred from DT40 into HPRT-deficient Chinese hamster ovary (CHO) cells by MMCT (**Figure 1A**). To monitor the HSA21 stability in cells, CAG promoter-driven EGFP was inserted into the loxP site of HSA21-loxP to generate the “HSA21-EGFP” chromosome through Cre-loxP recombination (**Extended Data Fig. 1A**). Recombinant clones were selected using HAT, and 6 out of 6 HAT-resistant clones were positive by PCR using Cre-loxP recombination-specific primers. The recombination process was confirmed by fluorescence in situ hybridization (FISH) (**Extended Data Fig. 1B-C**).

**Figure 1.**
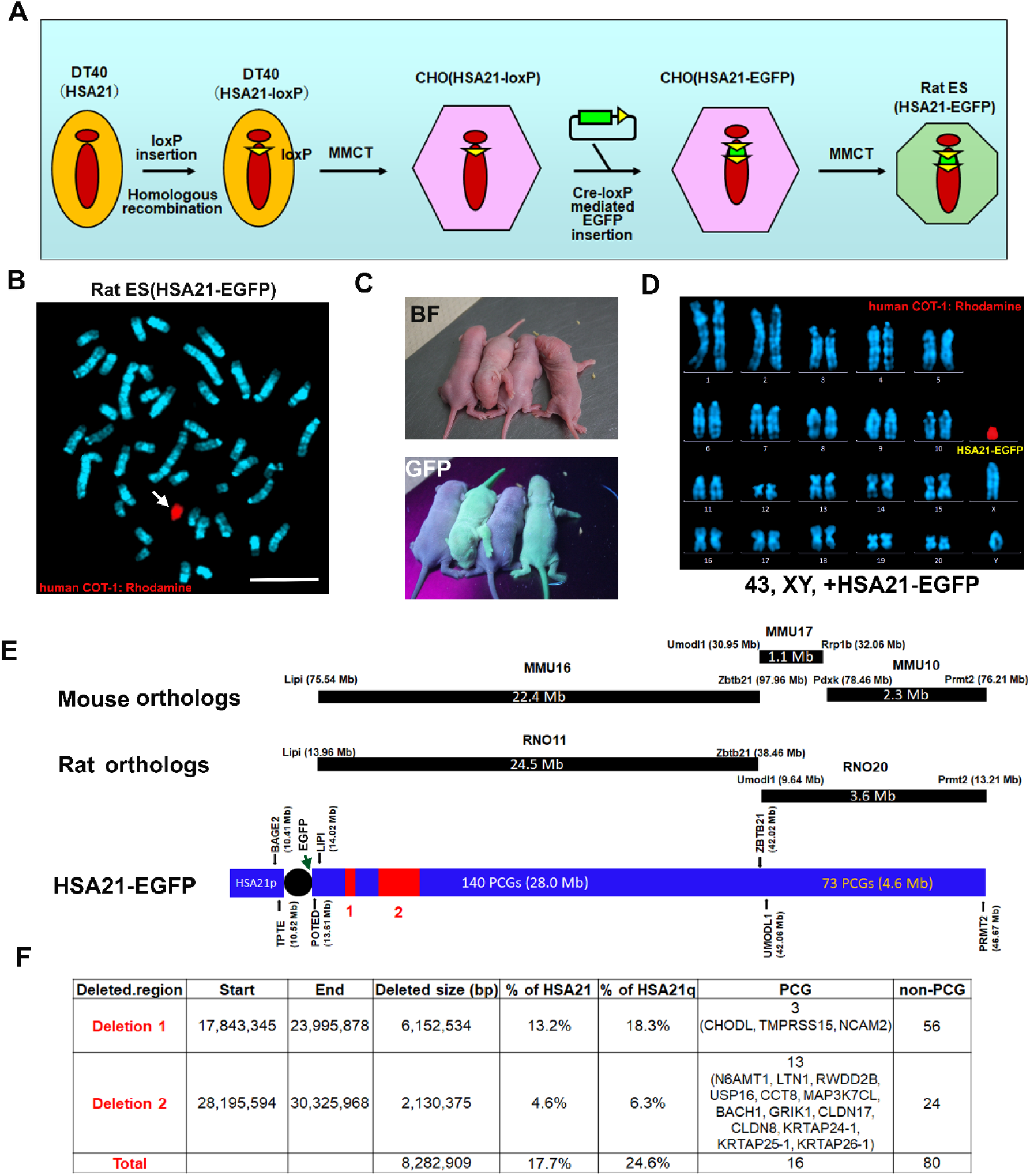
Generation of TcHSA21rat and WGS of the HSA21. (**A**) Schematic diagram of HSA21-EGFP construction and MMCT. (**B**) FISH of rat ES (HSA21-EGFP) cells. Digoxigenin (Rhodamine)-labeled human COT-1 DNA as the FISH probe for HSA21 and DNA counterstained with DAPI. Scale bar (10 m). (**C**) TcHSA21rat and euploid (Eu) pups visualized by GFP flashlights. (**D**) Karyotype of TcHSA21rat lymphocytes. Digoxigenin (Rhodamine)-labeled human COT-1 DNA as the FISH probe for HSA21-EGFP. (**E**) The orthologous relationships between HSA21, rat chromosomes, and mouse chromosomes. The HSA21-EGFP cartoon is normalized to PCG numbers. The positions and sizes of the two deletions detected by WGS are shown in red. (**F**) Summary of positions and genetic contents of the deletions in HSA21-EGFP.

HSA21-EGFP was then transferred from CHO cells into “BLK2i-1” rat embryonic stem (ES) cells (RGD ID: 10054010) via MMCT, producing two rat ES clones with a freely segregating HSA21-EGFP chromosome based on FISH analysis (**Figure 1B**). Cells from the rat ES clone with a major karyotype of 43,XY,+HSA21-EGFP were microinjected into the blastocoelic cavity of host Crlj:WI blastocysts (Charles River Laboratories Japan, INC., Kanagawa, Japan) to produce chimeras. Sixty-five offspring were produced from 125 rat ES-injected blastocysts, and 53 out of 65 offspring (82%) were chimeric (**Supplementary Table 1**). Among 19 male chimeras, 1 rat had a GFP-positive seminiferous tubule (**Extended Data Fig. 1D**). Round spermatids of the GFP-positive male chimeric rat were injected into rat ooplasm using round spermatid injection (ROSI), and 5 of 13 offspring were GFP-positive (**Supplementary Table 2**). Three GFP-positive females were then crossed with Crlj:WI (Wistar) males to establish the first transchromosomic rat model of DS named “TcHSA21rat”. Pups could be recognized using a UV flashlight (**Figure 1C**). The supernumerary HSA21 was visualized in lymphocytes by FISH (**Figure 1D**).

Although the annotation of HSA21p in GRCh38.p12 shows 17 protein coding genes (PCGs), 15 of them are not well annotated (**Supplementary Table 3A**), which are currently either considered as HSA21q paralogs or genome assembly errors^32, 33^. *BAGE2* and *TPTE* may be the only two bona fide PCGs on HSA21p. *BAGE2* has no reported mouse or rat orthologs, while *TPTE* has an annotated mouse ortholog on MMU10 but does not have any rat ortholog. Three-way comparison of HSA21q orthologous relationships based on genome assemblies of human (GRCh38.p12), rat (Rnor_6.0), and mouse (GRCm38.p6) document evolution between these species (**Figure 1E**). After excluding 49 keratin associated protein genes (*KRTAP*s), 158 out of 164 HSA21q PCGs have orthologs in both rat and mouse, the exceptions being *POTED, TCP10L, AP000295.1, AP000311.1, SMIM34A,* and *H2BFS*. The “*LIPI-ZBTB21*” orthologous segments are found on RNO11 and MMU16, while “*UMODL1-PRMT2*” orthologous segments are on RNO20 but are divided in mice between MMU17 and MMU10. The PCG arrangement in orthologous segments is the same among three species except for the rat *Hmgn1* (**Supplementary Table 3B**); this may be an annotation issue of the rat genome. WGS analysis showed two deletions in the HSA21-EGFP in TcHSA21rat (**Figure 1F** and **Supplementary Table 3C**). Deletion 1 (17,843,345 - 23,995,878) contains 3 PCGs, and the same deletion occurs in TcMAC21^15^. Deletion 2 (28,195,594-30,325,968) is specific to the TcHSA21rat and contains 3 KRTAPs and 10 non-KRTAP PCGs. Together, HSA21-EGFP of TcHSA21rat contains 214 out of 230 (93%) HSA21 PCGs and only misses 13 non-KRTAP PCGs.

### The presence of HSA21 causes global transcriptional imbalance in TcHSA21rat

Forebrain transcripts from three pairs of TcHSA21rat and euploid (Eu) littermates at postnatal day (P)1 were analyzed by RNA-Seq (**Supplementary Table 4A**). We reported our results in FPKM (fragments per kilobase of transcript per million mapped reads) and used FPKM=0.5 as the cutoff to determine whether the gene was expressed or not. The HSA21p PCGs, *BAGE2* (FPKM=7.9) and *TPTE* (FPKM=5.0), were expressed (**Supplementary Table 4B**). Among 213 HSA21q PCGs, all 49 KRTAPs were not expressed (<0.5 FPKM) as expected, and 76 of 164 (46.3%) non-KRTAP PCGs had >5 FPKM (**Figure 2A-B**). In contrast, 9 of 601 (1.5%) HSA21 non-PCGs had >5 FPKM. Neither HSA21 PCGs in deleted regions nor 5 PCGs with no mouse or rat orthologs (*TCP10L, AP000295.1, AP000311.1, SMIM34A,* and *H2BFS*) were expressed in the P1 forebrain.

**Figure 2.**
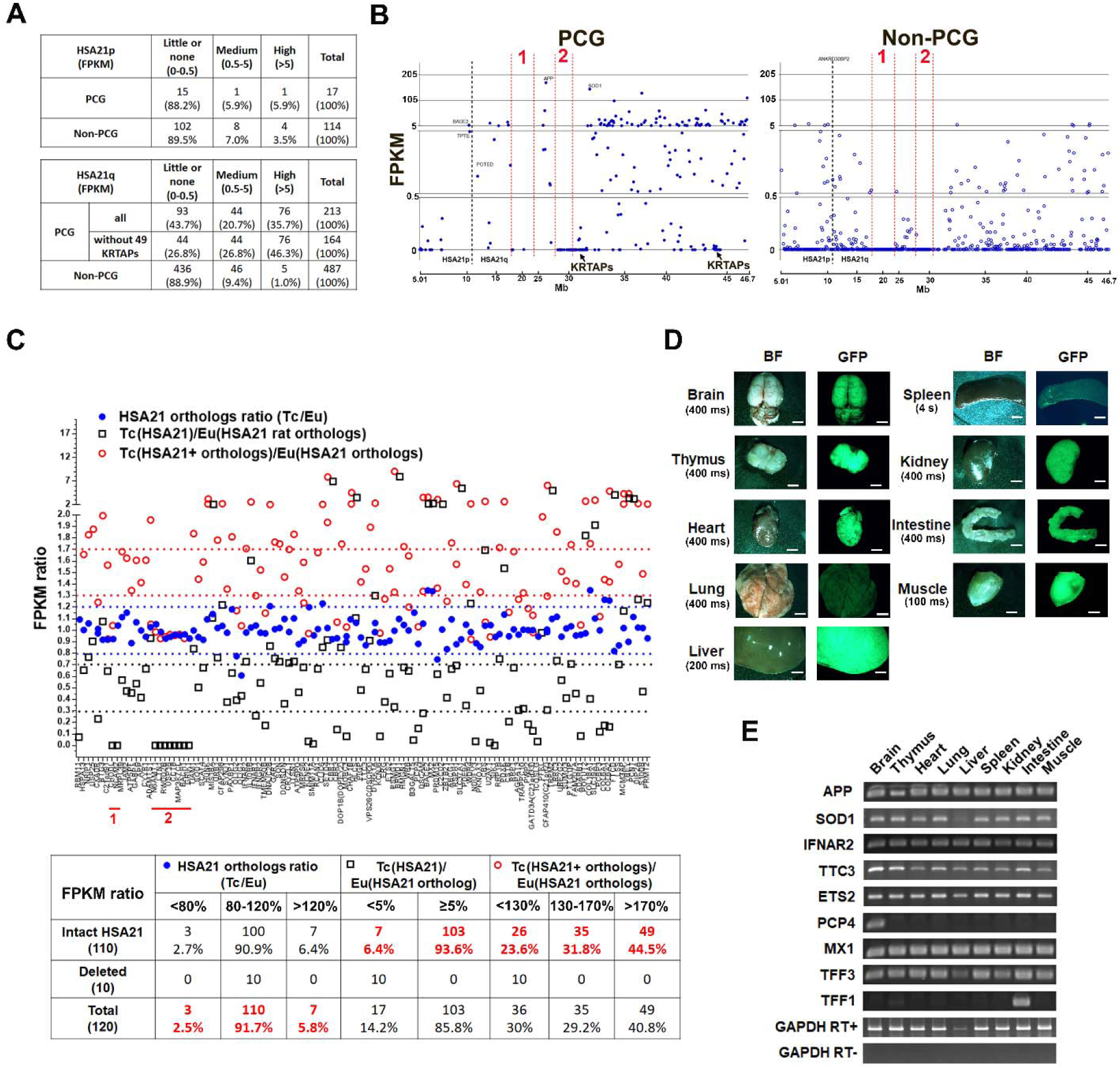
The presence of HSA21 causes global transcriptional imbalance in TcHSA21rat. (**A-C**) Forebrains of P1 male Eu and TcHSA21rat littermates analyzed by RNA-Seq (N=3 per group). (**A-B**) Transcription levels of HSA21 PCGs and non-PCGs in P1 forebrain of TcHSA21rat analyzed by RNA-Seq. (**C**) The comparisons of HSA21 PCGs and rat orthologs expression between TcHSA21rat and Eu among 120 HSA21 rat orthologs whose FPKM ≥1 in Eu. Three FPKM ratios are compared: Tc(HSA21)/Eu(HSA21 ortholog), the FPKM ratio of a HSA21 PCG in TcHSA21rat to its rat ortholog in Eu; HSA21 orthologs ratio (Tc/Eu), the FPKM ratio of a HSA21 rat ortholog in TcHSA21rat to that in Eu; Tc(HSA21+orthologs)/Eu(HSA21 orthologs), the FPKM ratio of total expression (sum of an HSA21 PCG and its rat ortholog) in TcHSA21rat to that in Eu. The PCGs in deletion 1 and 2 are indicated in red lines. Tc=TcHSA21rat. (**D**) TcHSA21rat organs visualized by GFP-UV light. The exposure time is shown, and BF=bright field, and scale bar (5 mm). (**E**) RT-PCR analysis of nine TcHSA21rat organs using 9 HSA21-specific primers.

The differential expression patterns of genes with FPKM ≥ 1 in the forebrain of P1 Eu rats were analyzed (**Supplementary Table 4C-D**). To examine if HSA21 affects the expression of HSA21 rat orthologs, 120 rat HSA21 orthologs with FPKM ≥ were compared between Eu and TcHSA21rat (**Figure 2C**): 110 were expressed at 80–120% of Eu levels in TcHSA21rat; 3 (*Olig1, Olig2,* and *Prdm15)* had reduced expression (<0.8 fold); 7 showed increased expression (>1.2 fold). To quantify total HSA21 overexpression levels in TcHSA21rat, the FPKM sum of an HSA21 PCG and its rat ortholog were compared between Eu and TcHSA21rat: 23.6% were in the low overexpression range (<1.3-fold increased over Eu); 31.8% in the expected range (1.3– 1.7-fold); 44.5% in the highly overexpressed range (>1.7-fold).

To analyze the HSA21 effect on global gene expression, we compared transcript levels of all rat genes (both non-coding and coding) between Eu and TcHSA21rat. More than 900 of 13473 rat genes with FPKM ≥ 1 in Eu were misregulated in trisomic rats (**Supplementary Table 5**): 358 were down-regulated (i.e., a rat gene in TcHSA21rat expressing less than 80% of Eu level); 570 were up-regulated (i.e., a rat gene in TcHSA21rat expressing more than 120% of Eu level). Human-specific RT-PCR for nine genes from RNA of nine organs showed that HSA21 genes were differentially expressed across TcHSA21rat tissues (**Figure 2D-E**). Together, these results indicate that in addition to altering HSA21 dosage, the presence of HSA21 causes global transcriptional imbalance in TcHSA21rat.

### The analysis of HSA21 mosaicism in TcHSA21rat

The transchromosomic mouse TcMAC21 shows stable transmission and retention of a hybrid chromosome containing HSA21q engineered with mouse centromere and telomere^15^. However, transchromosomic mice carrying human chromosomes or human artificial chromosomes (HAC) without mouse centromeres show high frequencies of mosaicism^34–36^, and retention rates of the foreign chromosome are lower in highly proliferative tissues, such as blood and spleen, than in slowly proliferative tissues like brain. Constitutive EGFP expression from HSA21-EGFP allows us to use flow cytometry (FCM) as a high-throughput tool to analyze HSA21 mosaicism. FCM of peripheral blood (PB) from 361 TcHSA21rats at 1-month-old or older showed 69.3% of TcHSA21rat with >90% GFP-positive rates and 81.4% of TcHSA21rats with >80% GFP-positive rates (**Figure 3A and Supplementary Table 6A**).

**Figure 3.**
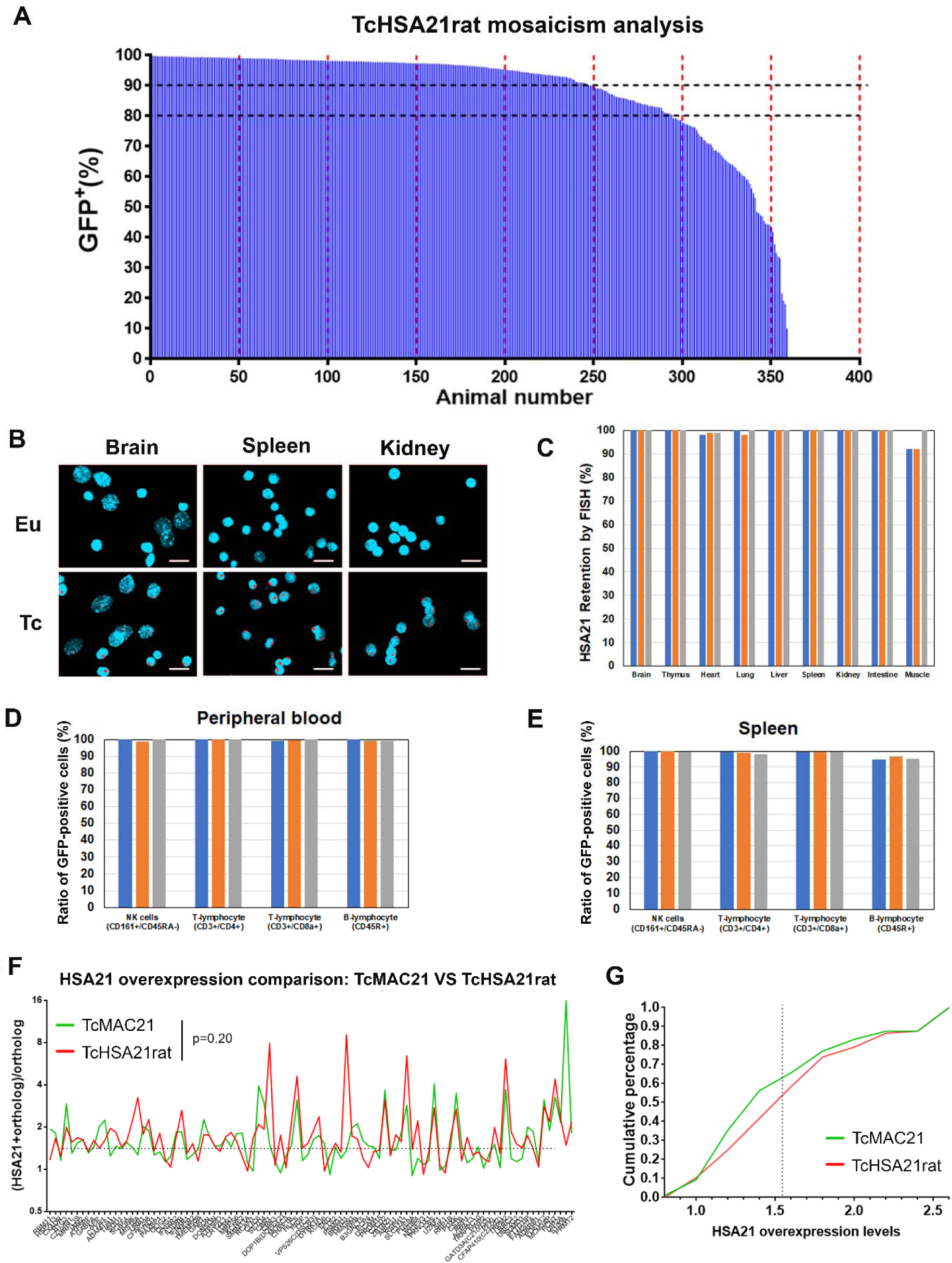
Mosaicism analysis of TcHSA21rat. (**A**) Mosaicism analysis of peripheral blood cells from 361 TcHSA21rats using FCM. (**B**) Representative FISH images of isolated cells from Eu and TcHSA21rat tissues using Digoxigenin (Rhodamine)-labeled human COT-1 DNA as the HSA21 probe. Scale bar (20 μm). (**C**) Summary of FISH of various TcHSA21rat tissues. Different rats are shown by different one-color bars, and n=3. (**D**) FCM of NK cells and lymphocytes from PB (n=3). (**E**) FCM of NK cells and lymphocytes from spleen (n=3). (**F-G**) The HSA21 overexpression pattern comparison between TcHSA21rat and TcMAC21 based on RNA-seq of P1 forebrain (Eu and TcHSA21rat littermates, n = 3 per group; Eu and TcMAC21 littermates, n = 3 per group). 96 HSA21 PCGs are compared based on two criteria: 1) HSA21 PCGs are intact in both TcMAC21 and TcHSA21rat, and 2) FPKM of HSA21 orthologs ≥ 1. (**F**) Overexpression pattern of individual PCG from TcHSA21rat and TcMAC21, and the differences are analyzed by Mann-Whitney test. (**G**) Cumulative distribution of overexpression levels. The overexpression levels > 2.5 are normalized to 2.5.

FISH of various tissues (brain, thymus, heart, lung, liver, spleen, kidney, small intestine, and skeletal muscle) from three TcHSA21rats with >90% GFP-positive rates in PB showed 92-100% HSA21 retention rates (**Figure 3B-C)**. FCM of NK cells (CD161^+^/CD45RA^-^), T-lymphocytes (CD3^+^/CD4^+^ or CD3^+^/CD8a^+^), and B-lymphocytes (CD45R^+^) from the same rats showed >95% GFP-positive rates (**Figure 3D-E**). Correlation analysis showed that HSA21 retention rates in PB measured by FISH were strongly correlated with GFP-positive rates measured by FCM (R^2^=0.9882, **Extended Data Fig. 2A**).

As TcMAC21 is not mosaic and contains almost the same HSA21 as TcHSA21rat, we compared HSA21 overexpression levels between TcHSA21rat and TcMAC21 based on RNA-seq data of P1 forebrain from three pairs of Eu and TcHSA21rat littermates and two pairs of Eu and TcMAC21 littermates (**Supplementary Table 6B**). Among HSA21 orthologs with FPKM ≥1 (116 in mice and 120 in rats), 113 of them were expressed in both models (**Extended Data Fig. 2B**). Ninty-six HSA21 PCGs were intact in both species and had FPKM ≥1 orthologs. HSA21 overexpression patterns were similar between the two DS models based on Mann-Whitney tests (p=0.2, **Figure 3F**) and cumulative frequency analysis (**Figure 3G**). Together, these results suggest that HSA21 mosaicism is minimal in TcHSA21rat and that non-mosaic TcHSA21rat could be easily screened.

### TcHSA21rat shows altered cognitive and behavioral phenotypes

At about three months of age, Eu and TcHSA21rat (>80% GFP-positive rates in PB, n=15 per group) males were assessed in the light/dark transition (LDT) test, the open field (OF) test, and the Morris water maze (MWM) test (**Figure 4A**). The LDT test, a widely used test to assess anxiety-like behavior in rodents^37, 38^, showed that both the time spent in the light compartment and the number of transitions between light and dark compartments were significantly decreased in TcHSA21rat compared with Eu (p<0.05, **Figure 4B-C**). In the OF test that has been widely used to evaluate novelty-induced exploratory activity in mice and rats^15, 39, 40^, TcHSA21rat showed a significant increase in the total distance traveled compared with Eu (p<0.01, **Figure 4E**).

**Figure 4.**
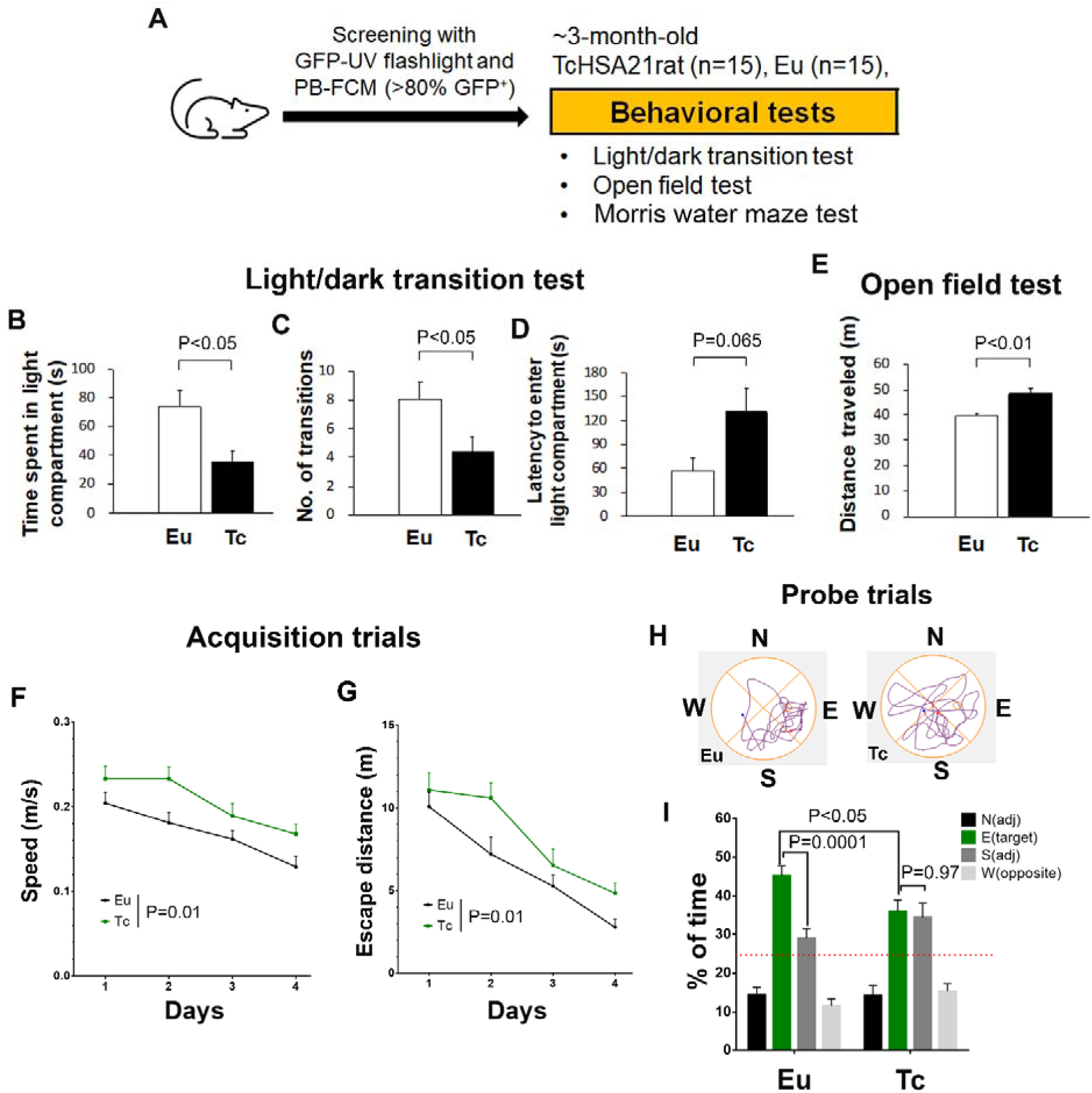
TcHSA21rat shows higher anxiety, hyperactivity, and learning and memory deficits. (**A**) The experimental design for behavioral tests (n=15 per group). (**B-D**) Light/dark transition test: (**B**) the time spent in the light compartment; (**C**) the number of transitions between light and dark compartments; (**D**) The latency time to enter the light compartment. Data are analyzed by Student’s t-test (B and C) and Mann-Whitney U test (D). (**E**) Open field test. The total distance traveled is compared using Welch’s t-test. (**F-H**) Morris water maze test: (**F-G**) Swimming speed and escape distance in acquisition trials, and data are analyzed by repeated measures two-way ANOVA; (**H**) Representative tracking plots of the probe trials; (**I**) probe trials, data are analyzed by two-way ANOVA, and Sidak’s post-hoc for comparing the % of time spent in each quadrant between Eu and TcHSA21rat, and Tukey’s post-hoc for analyzing whether animals prefer the target quadrant. All Data are expressed as mean with SEM. Eu=Euploid rat, Tc=TcHSA21rat.

The MWM paradigm consisted of four days of acquisition trials and a probe trial conducted 24 hours after the last acquisition trial was used to assess spatial learning and memory. As TcHSA21rat swam faster than Eu in acquisition trials (p=0.01, **Figure 4F**), the performance in acquisition trials was reported as escape distance rather than latency. Training improved escape distance in both groups, but the improvement was significantly slower in TcHSA21rat than Eu (p=0.01, **Figure 4G**). In probe trials, TcHSA21rat spent substantially less time than Eu in the target quadrant (p<0.05, **Figure 4H-I**). Unlike Eu that had a strong preference for the target quadrant (p=0.0001, target vs. adjacent), TcHSA21rat had no preference for the target quadrant (p=0.97, target vs. adjacent). Together, those data indicate that TcHSA21rat has higher anxiety, locomotor hyperactivity, and learning and memory deficits.

### TcHSA21rat has distinct alterations in brain morphology and cerebellar foliation

Brain morphometry of Eu and TcHSA21rat males at about four months of age (>80% GFP-positive rates in PB, n=9 per group) was analyzed using T2-weighted MRI. Total brain volume of TcHSA21rat was significantly smaller than Eu (2266 ± 80 mm^3^ vs. 2379 ± 73 mm^3^, p<0.005, **Figure 5A**). Among major brain structures, both neocortex (p=0.002) and cerebellum (p<0.0001) of TcHSA21rat were significantly smaller than those of Eu. The average cerebellar volume of TcHSA21rat was ∼86% of Eu. When normalized to the total brain, the percentage volume of cerebellum was disproportionately decreased in TcHSA21rat (14.1% in TcHSA21rat vs. 15.7% in Eu, **Figure 5B**). In contrast, the percentage volume of midbrain regions, such as thalamus and superior colliculus, were increased in TcHSA21rat.

**Figure 5.**
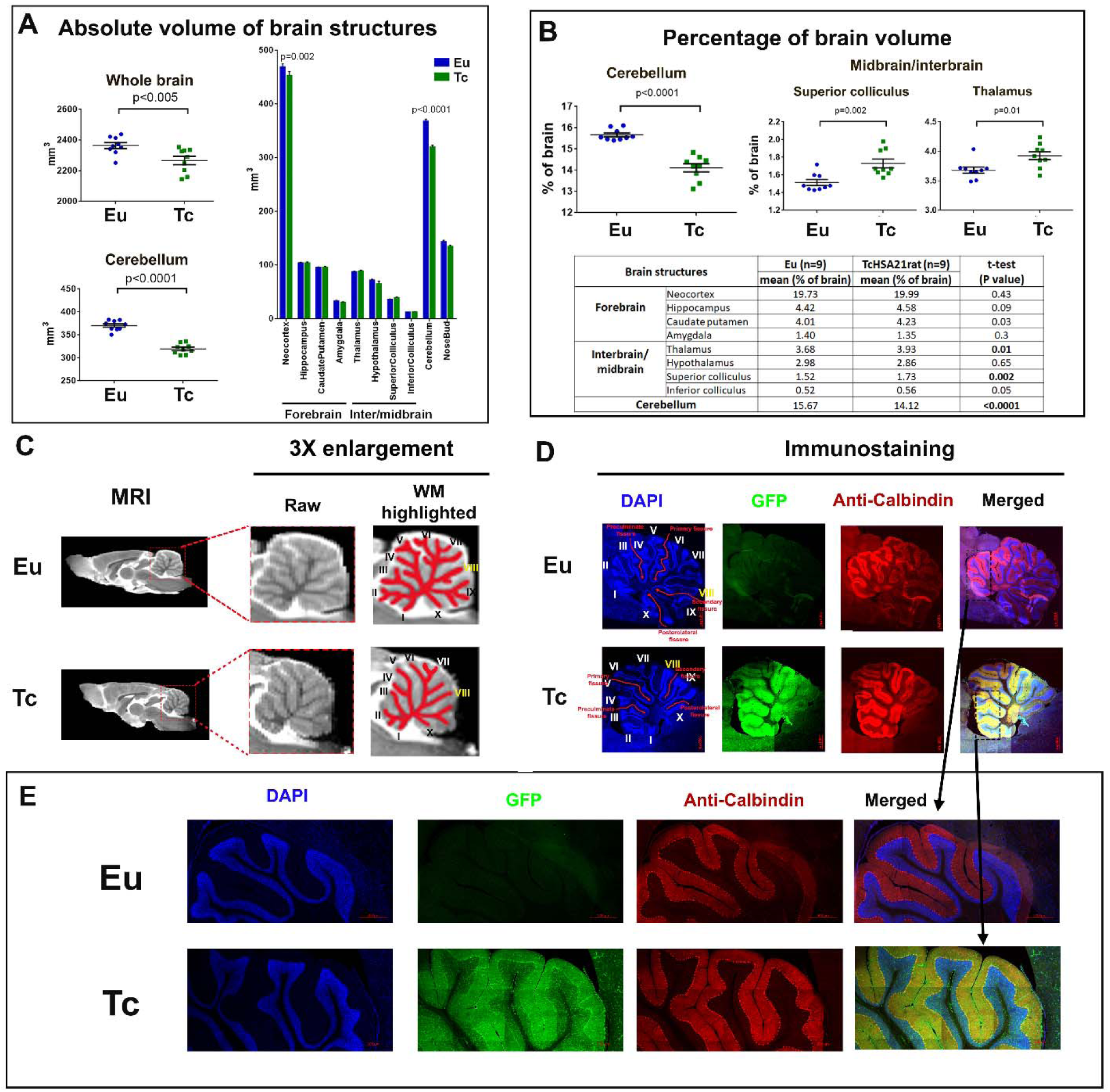
TcHSA21rat has distinct alterations in brain morphometry and cerebellar foliation. (**A**) Statistical analysis of absolute volumes of whole brain and its subregions based on T2-weighted MRI (n=9 per group). Data are analyzed by two-way ANOVA and Sidak’s multiple comparisons test. (**B**) Statistical analysis of percentage volume of brain subregions. Data are analyzed by unpaired t-tests. (**C**) Representative T2-weighted MRI of Eu and TcHSA21rat brains. Both raw cerebellum MRI and white matter (WM) highlighted MRI are shown. (**D-E**) Immunostaining of parasagittal brain sections with anti-calbindin antibody (red) and DAPI (blue). Eu=Euploid rat, Tc=TcHSA21rat.

Due to larger brain size, although the same MRI machine was used, there was better visibility of cerebellar vermian lobules (midsagittal section) in rat brain MRI **(Figure 5C**) than mouse brain MRI^15^. TcHSA21rat showed significantly fewer sublobules than Eu based on both MRI **(Figure 5C**) and histology analyses (**Figure 5D-E**). This phenotype was consistently observed in all nine TcHSA21rats analyzed (**Extended Data Fig. 3**). Together, these data show that TcHSA21rat not only recapitulates well-characterized DS brain phenotypes (i.e., smaller brain and disproportionately small cerebellum) but also has reduced cerebellar foliation, a feature not captured in DS mouse models.

### TcHSA21rat shows anomalies in craniofacial morphology, heart development, husbandry, and stature

Micro-computed tomography (micro-CT) images of the heads of about 4-month-old TcHSA21rat (n=7) and Eu (n=10) littermates were analyzed morphometrically using forty 3D cranial landmarks (lms) (**Figure 6A, Extended Data Fig. 4,** and **Supplementary Table 7**). Euclidean distance matrix analysis (EDMA)^41^ revealed statistically significant differences between TcHSA21rat and Eu for subsets of measures that defined the cranial base, cranial vault, and facial skeleton (p ≤ 0.025). The TcHSA21rat craniofacial skeleton is generally smaller than Eu. 90% confidence interval testing^42^ of linear distances between landmarks showed localized effects of HSA21 concentrated on the facial skeleton, with additional, more subtle effects on the cranial vault (**Figure 6A**). In agreement with the EDMA analysis, Generalized Procrustes based principal components analysis (PCA) of craniofacial shape variables revealed the facial skeleton of TcHSA21rat to be retracted and the posterior cranial vault to be “rounded” relative to Eu (**Figure 6B-C**). Principal component axis 1 (PC 1) explained 45.3% of the total variance in the sample and captured between-group differences between TcHSA21rat and Eu (**Figure 6B**). TcHSA21rat occupied the positive end on PC 1, indicating an antero-posteriorly retracted face associated with a supero-inferiorly “raised” cranial vault (**Figures 6A and 6C(i)**). PC 2 (12.7%) accounted for within-group variation, with TcHSA21rat showing more variation than Eu. Shape differences along this axis were mainly concentrated in the orientation of the snout and postero-inferior aspect of the neurocranium (**Figure 6C(ii)**).

**Figure 6.**
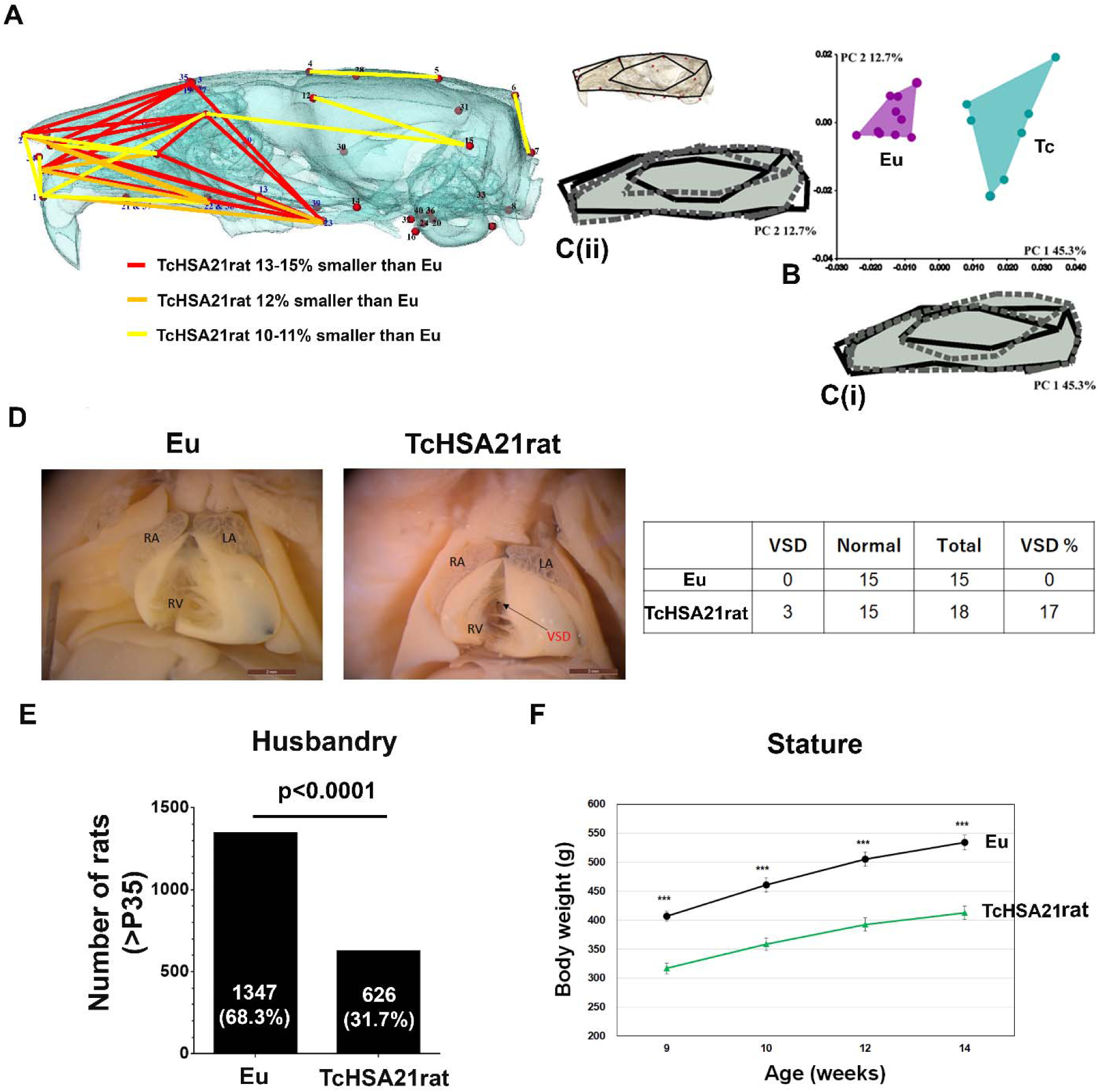
TcHSA21rat shows anomalies in craniofacial morphology, heart development, husbandry, and stature. (**A-C**) Craniofacial morphology analysis of Eu (n=10) and TcHSA21rat (n=7) in 4-month-old animals. (**A**) Statistically significant differences in cranial morphology between Eu and TcHSA21rat estimated by EDMA. The partially transparent bone segmented from HRμCT images is shown to better visualize the differences, with the rostrum at left and the occiput at right. The linear distances pictured are limited to those that differed significantly by ≥ 10% (using α = 0.10 confidence limits) and represent significantly smaller dimensions in TcHSA21rat. Different colored lines represent the percent differences. (**B**) PCA plot to show between-group differences (PC 1) and within-group variations (PC 2). (**C(i)**) Shape change on the positive end of PC 1 is represented by the dashed and shaded wireframe diagram of the cranium. Positive scores show a retraction of the snout and superiorly raised posterior cranial vault. The solid black wireframe represents the negative end of PC 1 and shows an elongated overall cranial shape. (**C(ii)**) Shape changes on PC 2 are mainly related to the snout and inferior cranial base orientation. The dashed and shaded wireframe and the solid black wireframe represent the shape changes on the positive end and the negative end of PC 2, respectively. (**D**) CHD analysis of Eu (n=15) and TcHSA21rat (n=18) at E20.5-E22.5 by wet dissection. Of 18 TcHSA21rat hearts, three hearts showed ventricular septal defect (VSD, arrow). LA, left atrium; RA, right atrium; RV, right ventricle. (**E**) The offspring number of TcHSA21rat mothers from 263 litters. Only P35 or older offspring are counted. Data are analyzed by both the Chi-square test (p<0.0001) and the Binomial test (p<0.0001). See supplementary table 8 for more details. (**F**) Body weight difference between TcHSA21rat and Eu males at 9,10,12, and 14 weeks old (n=15 per group). Data are analyzed by the Student’s t-test (***p<0.001).

The hearts of TcHSA21rats (n=18) and Eu (n=15) littermates were analyzed by wet dissection at E20.5-E22.5, showing that 3 (17%) TcHSA21rat hearts and none of Eu hearts had ventricular septal defect (VSD) (**Figure 6D**).

As no fertile TcHSA21rat males have been found, TcHSA21rat is maintained through female transmission. The breeding record of 263 litters showed a trisomy frequency of 32% among 1973 offspring at age P35 or older, with an average litter size of 7.5 (**Figure 6E**). Comparison of trisomy frequency at E20.5-E22.5 and at P35 (**Supplementary Table 8**) indicated a perinatal loss in TcHSA21rat. Mass of TcHSA21rat males measured from week 9 to week 14 was consistently lower than Eu counterparts, with an average weight reduction of ∼20% (**Figure 6F**).

## Discussion

This study has generated the first transchromosomic rat model of DS “TcHSA21rat” containing a freely segregating, EGFP-inserted HSA21 with 93% of the PCGs. HSA21 genes are expressed and cause an imbalance in global gene expression in TcHSA21rat. Young adult TcHSA21rat not only has learning and memory deficits but also exhibits robust DS brain phenotypes, including smaller brain volume, disproportionately small cerebellum, and reduced cerebellar foliation. TcHSA21rat has altered craniofacial skeleton, higher CHD prevalence, male infertility, higher postnatal mortality, and smaller stature. Our study provides the first validation of TcHSA21rat that could act as the “Pharma-preferred” model for DS research.

### Rat vs. mouse models for DS research

Rats lack the wealth of genetic stocks available for research in mice, although advances in both embryology and CRISPR have made the rat considerably more accessible as a genetic model. They possess several advantages for preclinical research compared to mice. The most obvious is size. Surgical interventions and assessments of anatomical changes are more accessible. A newborn rat is already 6-8 times the mass of a newborn mouse. Anatomical features are larger and as shown here for cerebellum, may be more elaborated. Behavior testing for rats is more nuanced than for mice, a positive for DS research. Finally, rats are the traditional model for pharmaceutical development, and this wealth of experience may pay a dividend in translation of preclinical results to drug trials.

The cerebral cortex of lissencephalic species, including mice and rats, is smooth^43^. However, cerebellums of both mice and rats are foliated into five cardinal lobes. Rats show more complex cerebellar foliation than mice. Compared with Eu counterparts, the nearly identical trisomy causes significantly reduced foliation in TcHSA21rat but not in TcMAC21 **(Extended Data Fig. 5)**. TcHSA21rat is the first DS animal model showing reduced cerebellar foliation. Moreover, the reductions in both the absolute and percentage volume in the TcHSA21rat cerebellum detected by MRI are more robust than observed in other aneuploid mouse models of DS (Ts65Dn^44^, Tc1^45^, and TcMAC21^15^). The larger cerebellar size facilitates assessment of cerebellar cortical changes in TcHSA21rat and, with attenuated foliation, should provide a useful model for understanding cerebellar hypoplasia and its role in DS cognitive phenotypes.

People with DS have smaller overall brain volume ^46^, which is not well recapitulated in adult (> 3-month-old) aneuploid DS mouse models^15, 44, 45^ but is readily observable in TcHSA21rat by MRI. As with mouse models, the changes in the rat comprise midface skeletal retrusion plus a reduced mandible, recapitulating changes in the DS craniofacial skeleton.

A major goal of DS research is to ameliorate the cognitive impact of trisomy. Both anxiety and attention deficit hyperactivity disorder (ADHD) are significantly more frequent in children with DS^47–50^. DS mouse models, Ts65Dn and Tc1, show hyperactivity but not anxiety-like behaviors^51–54^. Here, we demonstrate that in addition to learning and memory deficits, TcHSA21rat shows both anxiety-like behavior and hyperactivity.

### Mosaicism and TcHSA21rat

About 90-95% of people with DS are characterized as non-mosaic DS, while 2–4% have segmental trisomy for a portion of HSA21 due to duplication or translocation, and 2–5% have mosaic DS^55^. However, the frequency of mosaic and translocation DS is not known with precision. There is no rigorous standard for declaring an individual to have mosaic DS. Although some laboratories use a more stringent criterion, typically the occurrence of trisomy 21 in 16 or more of 20 metaphases from peripheral blood cells is considered to be non-mosaic DS^55^. Individuals with mosaic DS generally have a milder clinical presentation than non-mosaic DS during the prenatal and perinatal period; it is not unusual for mosaic DS to be detected in older children or adults^56, 57^. According to a population-based study by Devlin and Morrison^58^, only 37.5% of mosaic DS was detected by clinical examination, compared with a near 100% detection rate of non-mosaic DS.

Here, as HSA21 in both TcHSA21rat and TcMAC21 are marked by EGFP, we are able to determine mosaicism frequency in PB with high precision using FCM. About 70% of TcHSA21rats have >90% GFP-positive rate in peripheral blood cells, while about 80% of TcHSA21rats meet the usual laboratory definition of non-mosaic DS (i.e., >80% trisomic karyotypes). The previous examination of TcMAC21 mice shows little mosaicism^15^. That observation is extended here by PB-FCM of 363 trisomic TcMAC21 mice, which shows that ∼95% of TcMAC21 mice are not mosaic when using the GFP-positive rate of 80% as the cutoff (**Extended Data Fig. 6**). TcHSA21rat with a high percentage of trisomic PB showed similarly high HSA21 retention rates in different tissues as seen in the TcMAC21 mouse. Through TcHSA21rat female transmission, the average litter size is 7.5, and 32% are GFP-positive when measured at P35. Therefore, the average number of non-mosaic TcHSA21rat per litter is 1.7 (90% GFP-positive rate in PB-FCM as cutoff) or 2.0 (80% GFP-positive rate in PB-FCM as cutoff), similar to frequencies of some successful mouse DS models.

### Future research

Human chromosomes are not mitotically stable in mouse cells and are stochastically lost over time^31, 59–62^. Previous studies indicate that the loss rate per mitosis may vary from a few tenths of one percent to a few percent^59, 61, 63^. Prior analysis of mosaicism in hepatocytes from nine Tc1 transchromosomic mice suggests that the retention rates of HSA21 vary from 35%-75% in the liver, assuming similar efficiencies of RT-PCR amplification of human and mouse orthologs^36^. Similarly, transchromosomic mice containing an EGFP-labeled, HSA21-derived HAC (21HAC2) show < 50% GFP-positive rates by PB-FCM (**Extended Data Fig. 7**). In contrast to mice, the rat shows greater tolerance to human centromeres. Elucidating reasons for this difference may facilitate the production of humanized mouse or rat models.

Reduced cerebellar foliation is a feature not captured in previous DS mouse models. It will be interesting to use neuroimaging to validate reduced foliation/gyration in the DS brain. High-resolution MRI capable of detecting alterations at microstructure (cellular) levels could enhance the MRI application to monitor disease progression in DS or neurological disorders. We recently used oscillating gradient diffusion MRI to detect subtle changes in different cerebellar layers of Ts65Dn that are not detectable in conventional MRI^64^. TcHSA21rat is a useful model for generating a rat cerebellar microstructure atlas for diffusion MRI over the course of development.

TcHSA21rat shows robust neurological phenotypes, replicating and extending observations in DS mouse models. Testing current candidate approaches for therapies targeting neurological phenotypes of DS (genes, peptides, or small molecules) will help prioritize clinical trials. In this regard, it will be interesting to study the TcHSA21rat regarding Alzheimer’s disease (AD). All people with trisomy 21 develop AD-like histopathology by the fourth decade and despite multiple failures in contrast to the general population, where AD is not generally diagnosed until the onset of behavioral symptoms. Attempts to ameliorate the onset of AD symptoms remain largely focused on eliminating amyloid plaques, and TcHSA21rat might support or refine these attempts.

### Concluding remarks

TcHSA21rat meets the genetic criteria for a good DS model, including aneuploidy (freely segregating chromosome introducing an extra centromere), minimal mosaicism, a large number of functionally trisomic HSA21 genes/orthologs, and no functionally trisomic or monosomic non-HSA21 genes/orthologs. With robust DS-related phenotypes, TcHSA21rat has the potential to accelerate the preclinical studies and translation of DS basic research in the coming decade.

## Methods

### Generation of rat ES cell line with HSA21

The HSA21-EGFP was constructed using a previously described Cre-loxP mediated gene insertion system with the HSA21^65^. A loxP site was inserted at position 13,021,348-13,028,858, NC_000021.9 in a HSA21 in DT40 cells as described previously^30^. The modified HSA21 (HSA21-loxP) was transferred to CHO cells via MMCT as described previously^31^. HPRT-deficient CHO cells (CHO HPRT^−/−^) containing HSA21 were maintained in Ham’s F-12 nutrient mixture. A plasmid vector containing the EGFP flanked by HS4 insulator, loxP and 3’HPRT (I-EGFP-I-loxP-3’HPRT), and Cre-recombinase expression vectors were transfected into CHO cells containing the HSA21-loxP using Lipofectamine 2000. The cell culture and colony expansion were performed as described previously^30^. The site-specific EGFP insertion into the HSA21-loxP was confirmed by PCR and FISH analyses as described previously^30^. To generate the rat ES (HSA21-EGFP) cells, rat ES cells were fused with microcells prepared from the donor CHO hybrid cells containing the HSA21-EGFP and selected with G418 as described previously^65^. The rBLK2i-1 embryonic stem cells (RGD ID: 10054010, http://rgd.mcw.edu/wg/home) were used in this study.

### Generation of chimeric and Tc rat

Chimeric rats were generated by blastocyst injection of the rat ES (HSA21-EGFP) cells, as described previously^66^. Briefly, 8–10 cells were microinjected into the blastocoelic cavity of host Crlj:WI (Charles River Laboratories Japan, INC., Kanagawa, Japan) blastocysts. The re-blastulated embryos were transferred into the uteri of pseudopregnant Crlj:WI recipients to allow the full-term development to pups. The contribution of the ES cells in the resultant offspring was confirmed by their coat color and/or the GFP fluorescence. To examine the germline competency of the chimera to the F1 generation, round spermatid injection (ROSI)^67^ was applied with a few modifications. At 10-weeks-old, testes of chimeras were microdissected, and GFP-positive seminiferous tubules (**Extended Data Fig.1D**) were selected for ROSI. Round spermatids were FACS-sorted from the testicular cell suspension after a freeze-thaw procedure. Oocytes retrieved from superovulated Slc:SD (Japan SLC, Inc., Shizuoka, Japan) female rats at 4–5 weeks old were activated with 5 µM ionomycin in mR1ECM for 5 min and were cultured in mR1ECM for 40 min. Then, these oocytes were injected with the round spermatid followed by an additional culture in mR1ECM for 20 min and 5 µg/mL cycloheximide in mR1ECM for 4 h. The next morning, the ROSI oocytes were transferred into the oviducts of pseudopregnant Crlj:WI recipients and pups were examined for GFP expression.

### Animals

All procedures related to animal care and treatment were approved by each local University/Institutional Animal Care and Use Committee. TcHSA21rats and TcMAC21 mice were maintained on Wistar (Crlj:WI, Charles River) and BDF1 (C57BL/6J (B6) x DBA/2J (D2)), respectively. Rats and mice were maintained in a Tottori University animal facility with 12-hour light/12- hour dark cycle, temperatures of 20-26°C with 40-70% humidity, and fed with standard chow and in-cage automatic water.

ARRIVE guidelines (https://arriveguidelines.org/) were followed in the design and execution of the project. A consolidated table of demographic animal information for each experiment of each figure, including genetic background, age, gender, and sample size, is provided (**Supplementary Table 9**). Investigators were blind to sample genotypes in all assays. Investigators of behavior tests were blind to genotypes.

### FISH analyses

FISH was performed using fixed metaphase or interphase spreads. Slides were hybridized with digoxigenin-(or biotin-) labeled (Roche, Basel, Switzerland) human COT-1 DNA (ThermoFisher) to detect HSA21 and biotin-labeled I-EGFP-I-loxP-3’HPRT (ThermoFisher) to detect EGFP on the HSA21, essentially as described previously ^31, 68^.

### WGS

Tail DNA from two TcHSA21rats was purified and sequenced for two different runs using the Illumina NextSeq 500 system. A library prepared with TruSeq DNA PCR-Free LT Sample Prep Kit (Illumina, USA) was used for the first run, and a library prepared with Nextera Mate Pair Sample Prep Kit (Illumina) was used for the second run. After cleaning the sequence reads, the short reads were mapped to whole genome sequences of rat (NCBI Rnor_6.0) and human chromosome 21 (NCBI NC_000021.9) using CLC Genomics Workbench ver. 9.5. A total of 536 million reads were mapped to the reference genome. Among those reads, 3.8 million reads were mapped to HSA21, GRCh38.p13 Primary Assembly. Excluding zero coverage regions, the effective depth of coverage was 16.7×.

### RNA-Seq

RNA was extracted from forebrains of Eu and TcHSA21rat males (n=3 per group) at P1. Standard mRNA purification and library preparation were conducted using NEBNext® Poly (A) mRNA Magnetic Isolation (E7490) and NEBNext® Ultra™ II RNA Library Prep Kit (E7770). Library quality was assessed via Agilent 2100 Bioanalyzer for DNA High sensitivity DNA chip. The prepared library was sequenced using HiSeq2500 Flowcell with 100 bp paired-end reads, with each sample containing approximately 50-60 million reads. The sequences were assessed with fastqc, and 30bp were trimmed from each sequence to achieve higher accuracy. HSA21 reference was extracted and appended onto the whole rat genome reference sequence to create the modified reference. Reads were then aligned with TopHat2. Sim4 and Leaff were used for cross-species analysis. Standard DEseq methodology was used for differential gene expression analysis.

### Flow cytometry

To evaluate the percentage of GFP-expressing cells by FCM, peripheral blood cells were collected from TcMAC21 mice and TcHSA21rats through tail vein and treated with ammonium chloride solution (0.17M NH4Cl in distilled water) for hemolysis followed by FCM buffer (5% FBS / 1mM EDTA in PBS) substitution. Resuspended cells were filtrated with 40 μm pore cell strainers, and the levels of GFP expression were analyzed by Gallios (Beckman Coulter). Cells were illuminated with a 488-nm laser, and the fluorescence was detected using the FL1 525±40 nm bandpass filter. Peripheral blood mononuclear cells (PBMCs) were gated for analysis, and minimal 20,000 events in the gate were analyzed. The same GFP-positive gating was used for each group to differentiate GFP-positive and negative cells. The percentage of GFP-positive (GFP^+^) cells was calculated by the number of GFP-positive cells within PBMCs in the gate.

To evaluate the percentage of GFP-expressing cells in lymphocyte subsets, peripheral blood cells and spleen were collected from euthanized TcHSA21rats with isoflurane. Peripheral blood cells were stained with cell type specific antibodies, and additional samples were prepared by stained with corresponding isotype control antibodies. Cells were hemolyzed, washed with FCM buffer, and filtered with cell strainers for FCM analysis. The spleen was physically processed by mashing between the frosted ends of two glass slides, and dissociated cells were resuspended with 10 mL of FCM buffer and filtered with 40μm pore cell strainers. Cells were hemolyzed, resuspended with 10 mL of FCM buffer, filtered by 40μm pore cell strainers, and counted. For staining, 1×10^6^ cells were applied to each reaction and incubated on ice for 30 min, followed by washing with FCM buffer and filtration. Antibodies used in this assay were, mouse anti-rat CD45RA (OX-33, Biolegend), CD161 (10/78, eBioscience), CD3 (1F4, Biolegend), CD4 (W3/25, Biolegend), CD8a (OX-8, Biolegend) and CD45R (HIS24, BD Pharmingen) conjugated with PE, PerCP-eFluor710, Alexa Fluor 647, PE, PerCP and PE, respectively. FCM analysis was performed with Gallios using different combinations of laser (488-nm and 638-nm) and bandpass filter (FL1 525±40, FL2 575±30, FL4 695±30 for PerCP-eFluor710 or 675±10 for PerCP, and FL6 660±20 nm). The lymphocyte population was gated, and > 13,000 cells were analyzed. Positive and negative populations were determined with samples stained by isotype controls. The percentage of GFP-positive cells in each lymphocyte subset was calculated by the number of GFP-positive cells within the subset.

### Behavioral analyses

#### Experimental design

Rats used for behavioral tests were male F6-8 TcHSA21rat on Wistar background. Blood cells of GFP-positive and GFP-negative littermates were collected from tail veins at 3-5 weeks of age, and GFP-positive cell rates of PBMCs were analyzed by flow cytometry. TcHSA21Rats with >80% GFP-positive rates (TcHSA21rat, n=15), and GFP-negative littermates (Eu rats, n=15) were used. Each rat was handled by experimenters for 5 minutes at 9 weeks of age. The same rat was tested in light/dark transition test, open field test, and Morris water maze test at 10-13 weeks of age. Rats were placed in the behavioral testing room at least one hour before the start of tests.

#### Light/dark transition test

The light/dark box with grid floor (MELQUEST Co., Ltd., Toyama, Japan) was used in this test. The apparatus consisted of two transparent compartments (45 x 27 x 26 cm) divided by a black guillotine door, and it had black and transparent lids. Inside of one compartment was covered with black walls and floor plates. Another floor was covered with a white floor plate. The light intensity of the center of the light compartment was 200-201 lux. Each rat was placed in the dark compartment of the box with the guillotine door closed. Immediately after opening the door, the behavior of each rat was recorded for 5 min using an animal movement analysis system with infrared sensors (SCANET MV-40, MELQUEST Co., Ltd.). The time spent in the light compartment, the number of transitions between light and dark compartments, and the latency to enter the light compartment for the first time were analyzed.

#### Open field test

The gray square open field apparatus (70 x 70 x 40 cm, MUROMACHI Kikai Co., Tokyo, Japan) was used in this test. The apparatus was illuminated by indirect lighting, and the light intensity of the center of the field was 30 lux. Each rat was placed at the corner of the field, and the behavior of each rat was recorded for 10 min using ANY-maze Video Tracking System (Stoelting Co., Wood Dale, IL, USA). The distance traveled was analyzed.

#### Morris water maze test

The gray circle water pool apparatus (1.5 m in diameter, MUROMACHI Kikai Co.) with the water temperature at 23-25°C was illuminated by indirect lighting. The light intensity of the center of the pool was 21-24 lux. Visual cues surrounded the pool were the same during four training days and one probe test day. A transparent platform (12 cm in diameter, MUROMACHI Kikai Co.) was located at the same position in the target quadrant (E) and hidden 2 cm below the water surface during training days. Each training day had 4 trials, and the time interval between each trial was more than an hour. The first trial of the next day was started more than 24 hours after the first trial of the previous day. For learning the location of the platform, rats swam until they reached and stayed on it for 5 seconds. If rats couldn’t reach the platform within 60 seconds, an experimenter guided rats to the platform, and rats were allowed to stay on the platform for 30 seconds. The probe test was performed > 24 hours after the final trial of the fourth training day. Rats swam in the pool without the platform for 60 seconds. The behavior of each rat was recorded using ANY-maze Video Tracking System. The escape distance during training days and the percent of time spent in each quadrant in the probe test were analyzed. Three problematic trial data (one trial for each rat) caused by the tracking error were excluded.

### Brain morphometry by MRI

TcHSA21Rats with >80% GFP-positive rates in PB-FCM and Eu littermates at ∼ 4-month-old were used for the *ex vivo* MRI analysis (n=9 per group). Rats were perfused with PBS and then 4% PFA. Heads were post-fixed in 4% PFA at 4 °C for 1 week, then kept in PBS for 3 days. Heads were stored in Fomblin (Fomblin Profludropolyether, Ausimont, Thorofare, NJ, USA) to prevent dehydration during imaging in an 11.7 Tesla scanner (vertical bore, Bruker Biospin, Billerica, MA). 3D T2-weighted images were acquired with the resolution = 0.08 mm x 0.08 mm x 0.08 mm. For analysis, *ex vivo* images were first aligned to the template image using automated image registration software (Diffeomap, www.mristudio.org) and adjusted to an isotropic resolution of 0.125 mm × 0.125 mm × 0.25mm. We quantitatively measured volume from different brain structures by combining automated and manual editing of images in ROIEditor (www.mristudio.org).

### Craniofacial morphology by micro-CT

High-resolution µCT images with a voxel size of 0.0300 mm were acquired by the Center for Quantitative X-Ray Imaging at the Pennsylvania State University (www.cqi.psu.edu) using the HD-600 OMNI-X high-resolution X-ray computed tomography system (BioImaging Research Inc.). A minimum threshold of 70-100 mg/cm^3^ partial density HA (based on HA phantoms imaged with specimens) was used to reconstruct isosurfaces in Avizo Lite 9.7 (Visualization Sciences Group). Micro-CT images of the heads of TcHSA21rats with >80% GFP-positive rates in PB-FCM (n=7) and Eu (n=10) littermates at ∼ 4-month-old were analyzed morphometrically. 3D coordinates of forty biologically relevant landmarks were collected from the isosurfaces to use in analyses. Specimens were digitized three times, and intra-observer measurement error was minimized by averaging coordinates of the three trials (maximum accepted error in landmark placement=0.05 mm). Euclidean Distance Matrix Analysis (EDMA) was used to statistically evaluate skull shape differences by hypothesis test and confidence interval estimation^41^. EDMA is a 3D morphometric technique invariant to the transformation group, including translation, rotation, and reflection^42, 69^. Briefly, the original 3D coordinates of landmark locations are rewritten and analyzed as a matrix of all unique linear distances among landmarks called the form matrix (FM). An average FM is estimated for each sample^42^. The difference between samples is evaluated by estimating ratios of like linear distances using sample-specific average FMs. The resulting matrix of ratios, the form difference matrix (FDM), is a collection of relative differences among landmarks used to define the forms. A non-parametric bootstrap procedure (100,000 resamples) is used to obtain confidence intervals for elements (each corresponding to a linear distance) of the FDM^41^ that reveals the localized effects of HSA21 on the skull. We also include a non-parametric bootstrap assessment of the null hypothesis that the mean forms of two samples are the same^41^. We measure form differences of the facial skeleton, the cranial base, and the cranial vault using landmarks specific to those regions and the software WinEDMA ( https://getahead.la.psu.edu/resources/edma).

Generalized Procrustes Analysis (GPA) was used to extract shape coordinates from the landmarks measured on the entire craniofacial skeleton. GPA calculates shape coordinates from the original landmark dataset by translating, scaling and rotating the data, and subsequently yielding Procrustes shape coordinates. A measure of overall size, called Centroid size (CS), is calculated as the square root of the sum of squared Euclidean distances from a set of landmarks to their centroid^70, 71^. Using the Procrustes coordinates, patterns of shape variation in the dataset were explored using principal component analyses (PCA). PCA is based on an eigenvalue decomposition of a covariance matrix, transforming Procrustes shape coordinates into scores along principal components (Slice 2007). PCA was performed using all forty cranial landmarks in our dataset. All other morphometric and statistical analyses, including the construction of the wireframe diagrams, were performed in MorphoJ (Klingenberg lab) and R programming software (version 4.0.1).

### Immunostaining

After MRI scan, brains of TcHSA21rat and Eu (n=4 per group) were removed, fixed in 4% PFA overnight at 4°C, and transferred to 30% sucrose for 48h. 30 or 40 μ cryosections were collected onto glass slides. Dry slides were post-fixed in 4% PFA and treated with 0.5% Triton in PBS. Parasagittal sections were immunostained with anti-calbindin (Cell Signaling, #13176) and DAPI.

### CHD analysis by wet dissection

E20.5-22.5 rat fetuses were removed and sacrificed, and hearts were flushed with PBS via the umbilical vein and then fixed in 4% PFA. The hearts were examined for cardiovascular anomalies under a dissecting microscope.

### Statistical Analysis

For each experiment, we stated statistical information, including the exact sample size, statistical tests, and the exact p-values in each figure or its legend. Unless otherwise noted, data were expressed as mean ± SEM (the standard error of the mean), and p value<0.05 was considered as statistically significant. Statistical programs including Microsoft Excel (Microsoft, USA), SPSS Statistics 25 (IBM, USA), GraphPad Prism (GraphPad Software, USA), and R programming software (version 4.0.1) were used.

### Data availability

All raw read data of WGS were deposited to DDBJ Sequence Read Archive (DRA) under accession number DRA010895.

## Acknowledgments

We thank Toko Kurosaki, Yukako Sumida, Masami Morimura, Kei Yoshida, Eri Kaneda, Akiko Ashiba, Megumi Hirose, Dr. Kazuomi Nakamura, Dr. Takashi Takeuchi, Manami Oka, Rina Ohnishi, and Michika Fukino at Tottori University, and Kayoko Morimoto at Trans Chromosomics Inc., for their technical assistance; as well as Dr. Hiroyuki Kugoh, Dr. Tetsuya Ohbayashi, Dr. Hiroyuki Satofuka, Dr. Takashi Moriwaki and Dr. Takahito Ohira at Tottori University and Dr. Oomiya Yoshihiro at Health Research Institute, National Institute of Advanced Industrial Science and Technology (AIST) for critical discussions. This research was partly performed at the Tottori Bio Frontier managed by Tottori prefecture. The work was supported in part by JST CREST Grant Number JPMJCR18S4, Japan (Y.K.), a grant of General Collaborative Project from the National Institute for Physiological Sciences, Japan (NIPS; 17-252 to Y.K.), The Mitsubishi foundation (M.O.), R01HD038384 (R.H.R), R21HD098540 (R.H.R.).

## Author contributions

Y.K., F.J.G., and R.H.R. conceived of the project; Y.K., F.J.G., M.Y., M.Hirabayashi, K.K., N.K., S.M.T., S.A., M.S., H.H., H.K., S.I.,Y.H., M.K., H.T., S.T., Y.N., M.Hiratsuka, Y.I., S.M., N.N., Y.L., A.J.M., B.C., and N.S. performed and analyzed all remaining experiments; J.T.R., M.O., and R.H.R. designed experiments and provided feedback during the project; Y.K. and F.J.G wrote the paper and prepared the figures. All authors commented and approved the manuscript.

## Materials & Correspondence

Correspondence and requests for materials should be addressed to Roger Reeves or Yasuhiro Kazuki.

## Additional information

### Extended Data

This file also contains 7 Extended Data Figs that are shown below.

### Supplementary information

The paper contains 9 Supplementary Tables, which were uploaded as separate files.

---

**Extended Data Fig.1.**
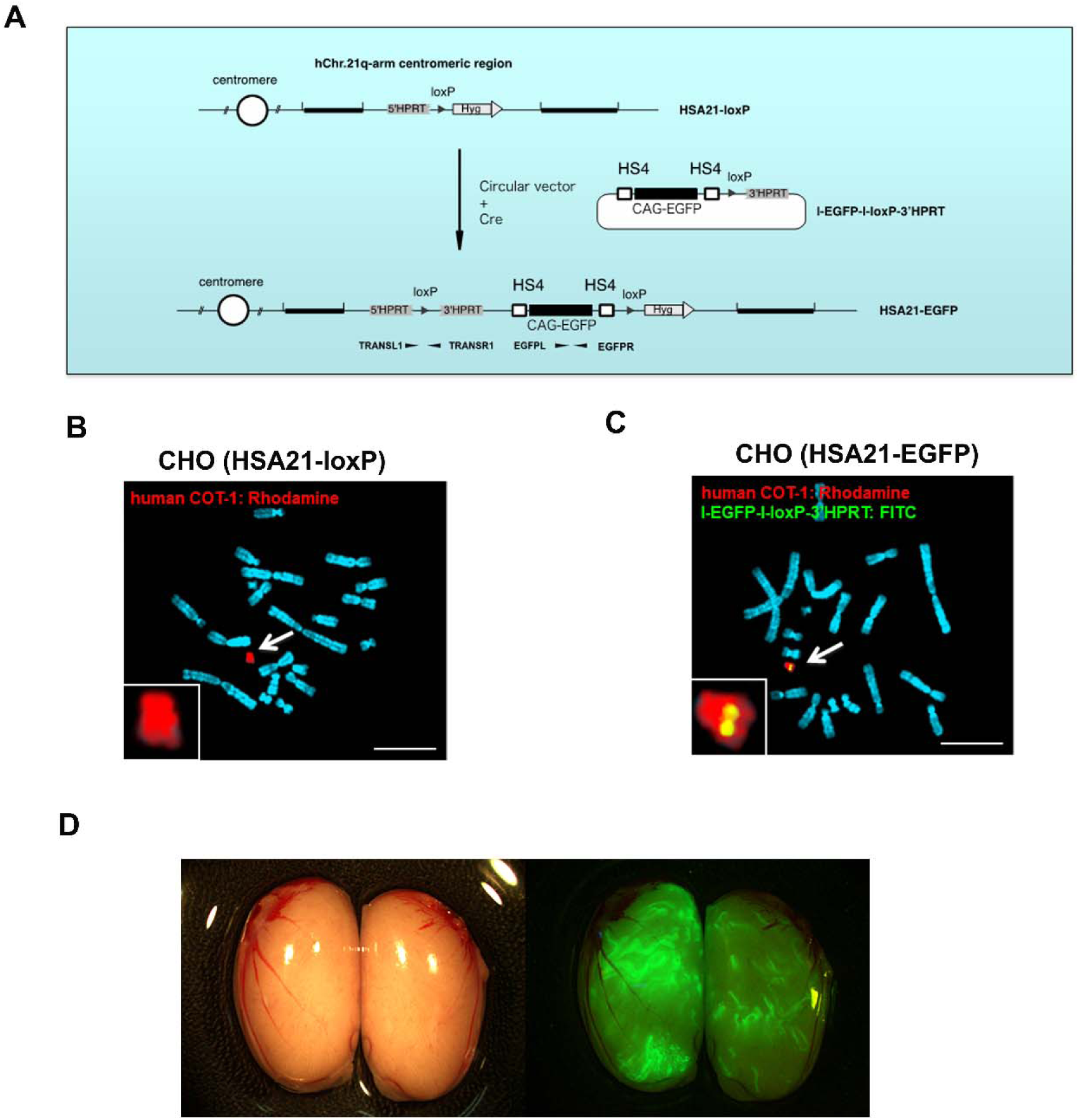
Generation of TcHSA21rat. (**A-C**) HSA21-EGFP chromosome generation through Cre-loxP recombination: (**A**) The information of vectors; (**B**) before Cre-loxP mediated recombination, CHO cells containing an HSA21-loxP; Rhodamine-labeled human COT-1 DNA as the FISH probe for HSA21-loxP; (**C**) after recombination, CHO cells containing the HSA21-EGFP; both Rhodamine-labeled human COT-1 DNA and FITC-labeled I-EGFP-I-loxP-3’HPRT probe used as FISH probes for HSA21-EGFP. Scale bar (10 μm). (**D**) GFP-positive seminiferous tubules in testes of chimera at 10 weeks old.

**Extended Data Fig.2.**
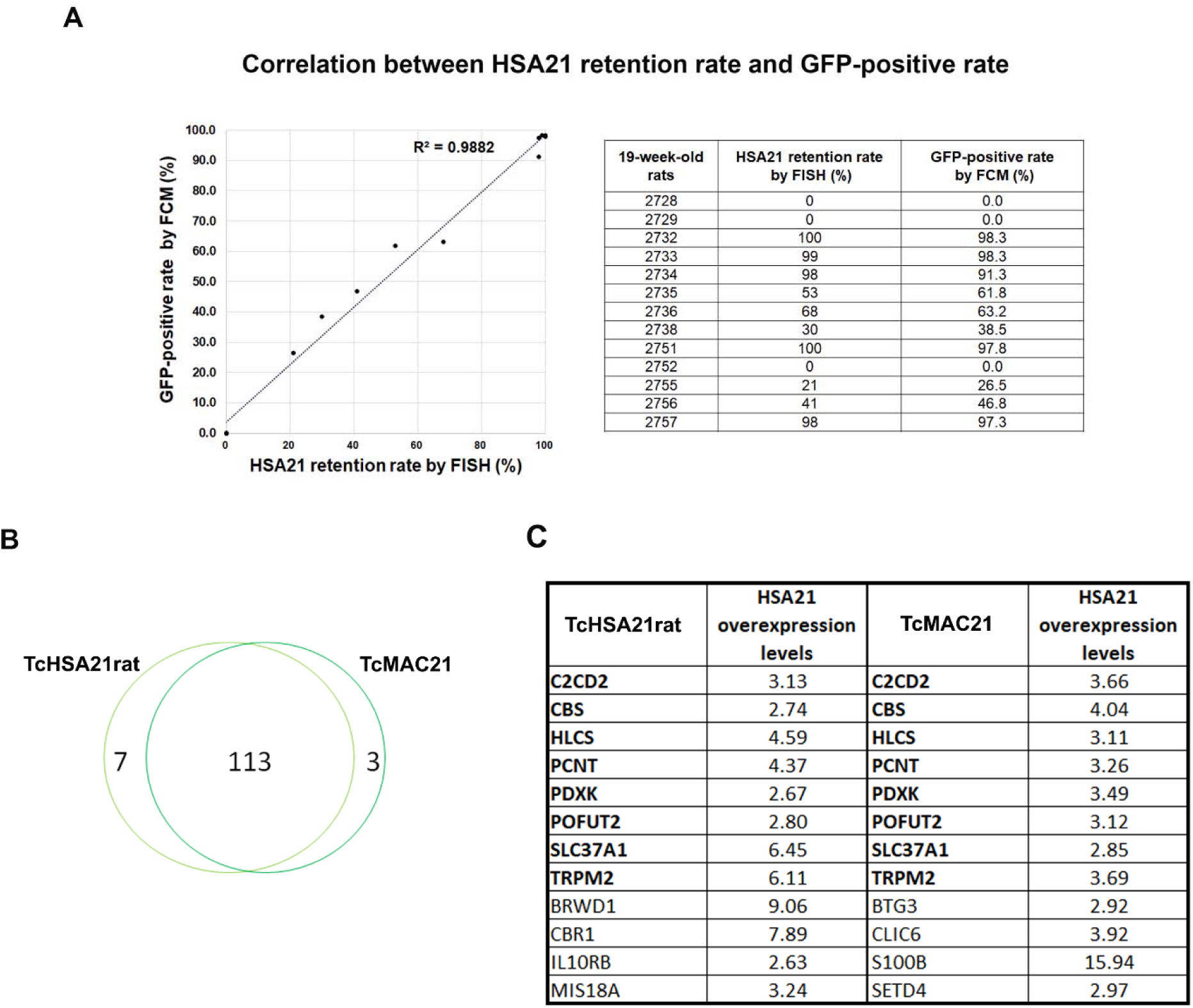
Mosaicism analysis. **(A)** The correlation analysis between the HSA21 retention rate measured by FISH and the GFP-positive rate measured by FCM in PB. **(B)** The HSA21 overexpression pattern of TcHSA21rat and TcMAC21 is based on RNA-seq of P1 forebrain. 116 in TcMAC21 and 120 TcHSA21rat have HSA21 ortholog FPKM ≥1, and 113 of them are common ones. (**C**) The list of HSA21 PCGs that are highly overexpressed. The cutoff of highly overexpressed is that the ratio of Tc (HSA21+ its ortholog) to Eu (HSA21 ortholog) is above 2.5. Eight of 12 are the same in TcHSA21rat and TcMAC21.

**Extended Data Fig.3.**
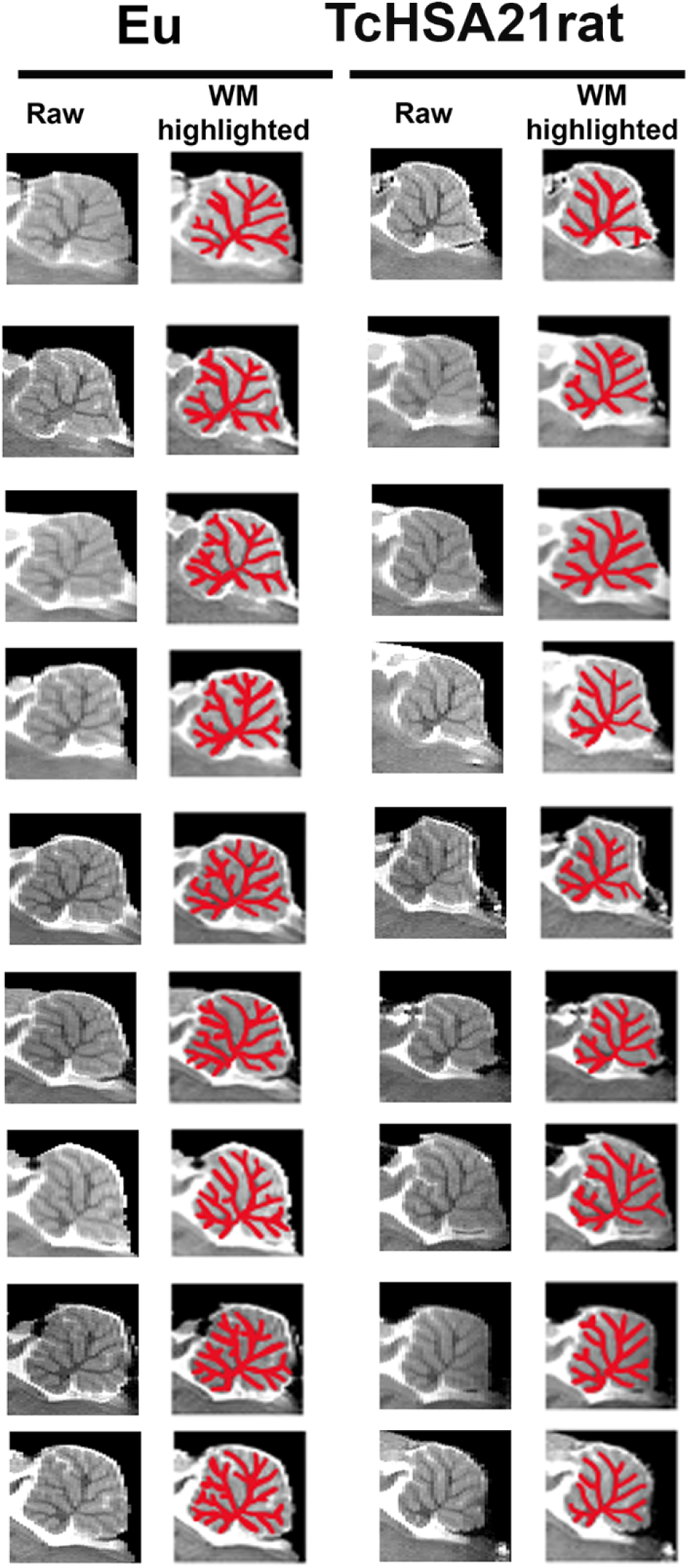
All MR images and related white matter highlighted images of cerebellums are shown.

**Extended Data Fig.4.**
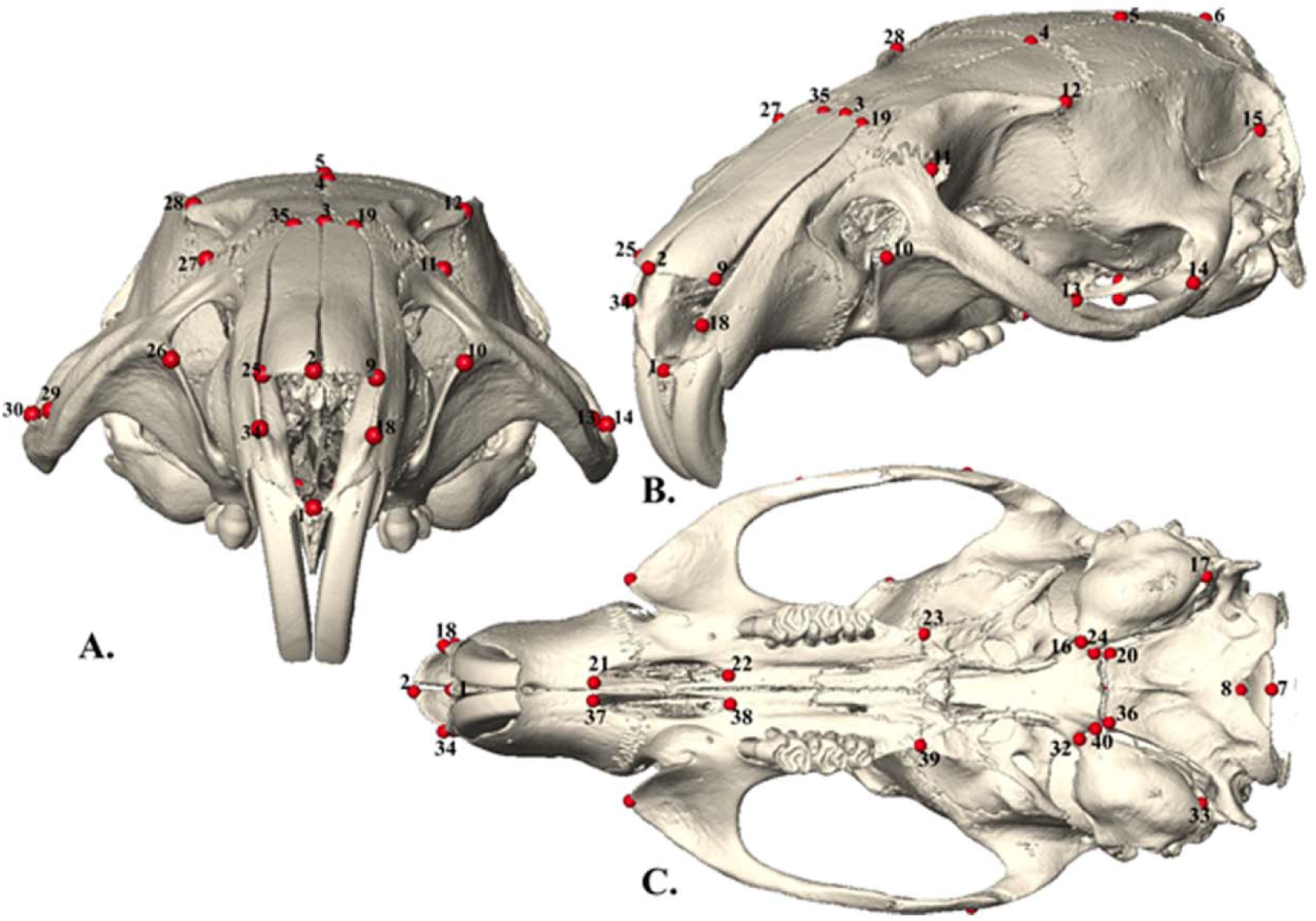
**(A)** Anterior view of rat cranium with landmark numbers. **(B)** Lateral oblique view of rat cranium with landmark numbers (lms #31 not shown). **(C)** Inferior view of rat cranium with landmark numbers.

**Extended Data Fig.5.**
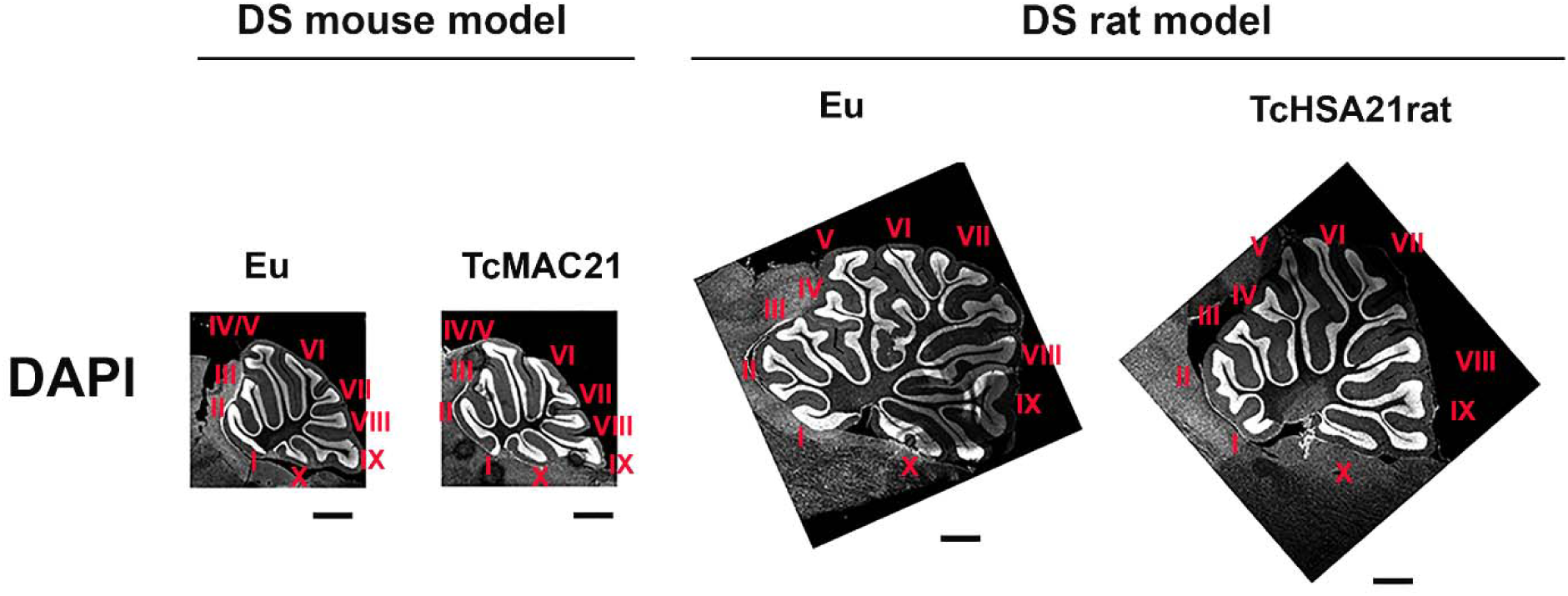
Cerebellar foliation analysis of the DS mouse model TcMAC21 and the DS rat model TcHSA21rat visualized by DAPI staining.

**Extended Data Fig.6.**
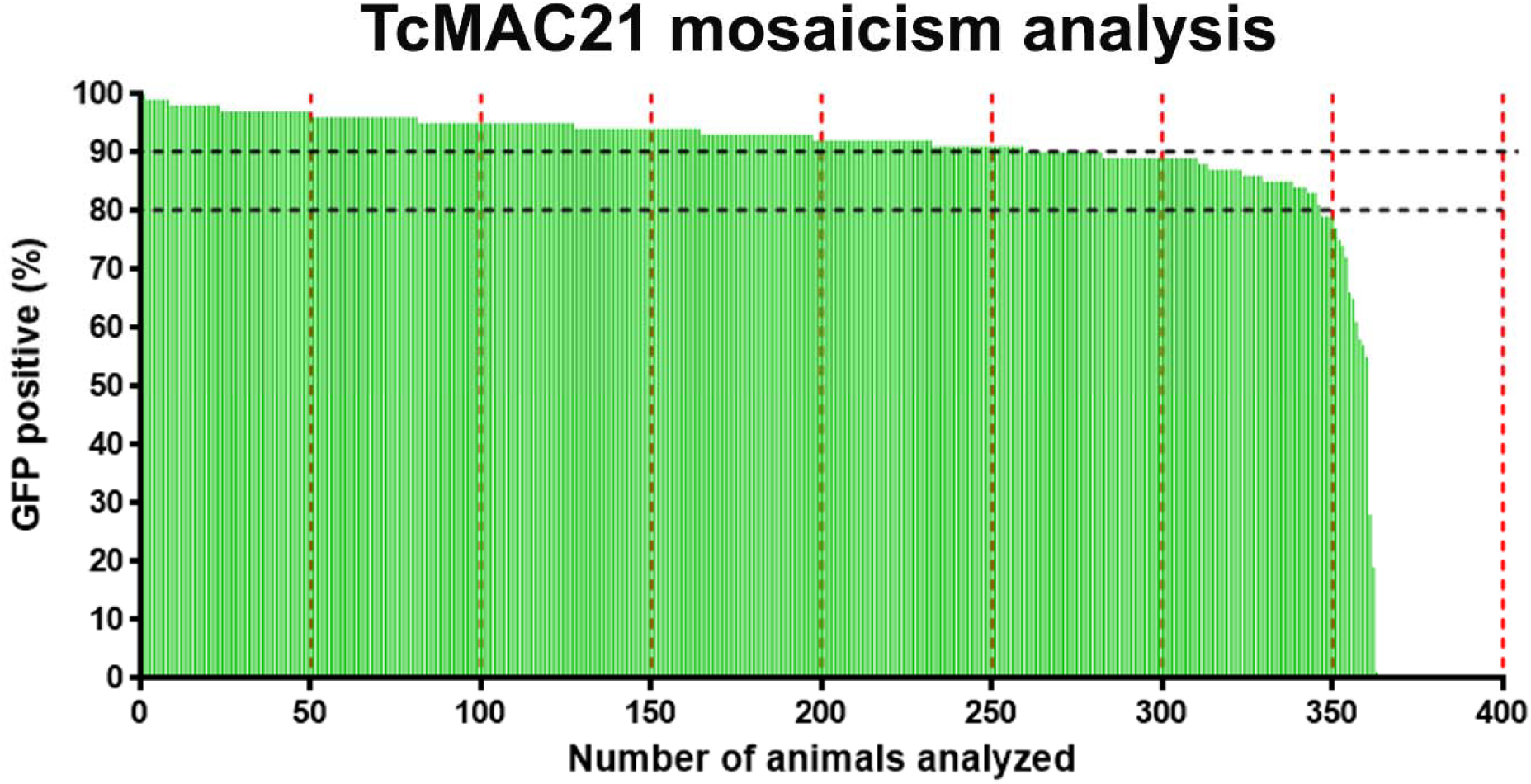
Mosaicism analysis of peripheral blood cells from TcMAC21 mice (n=363) using FCM.

**Extended Data Fig.7.**
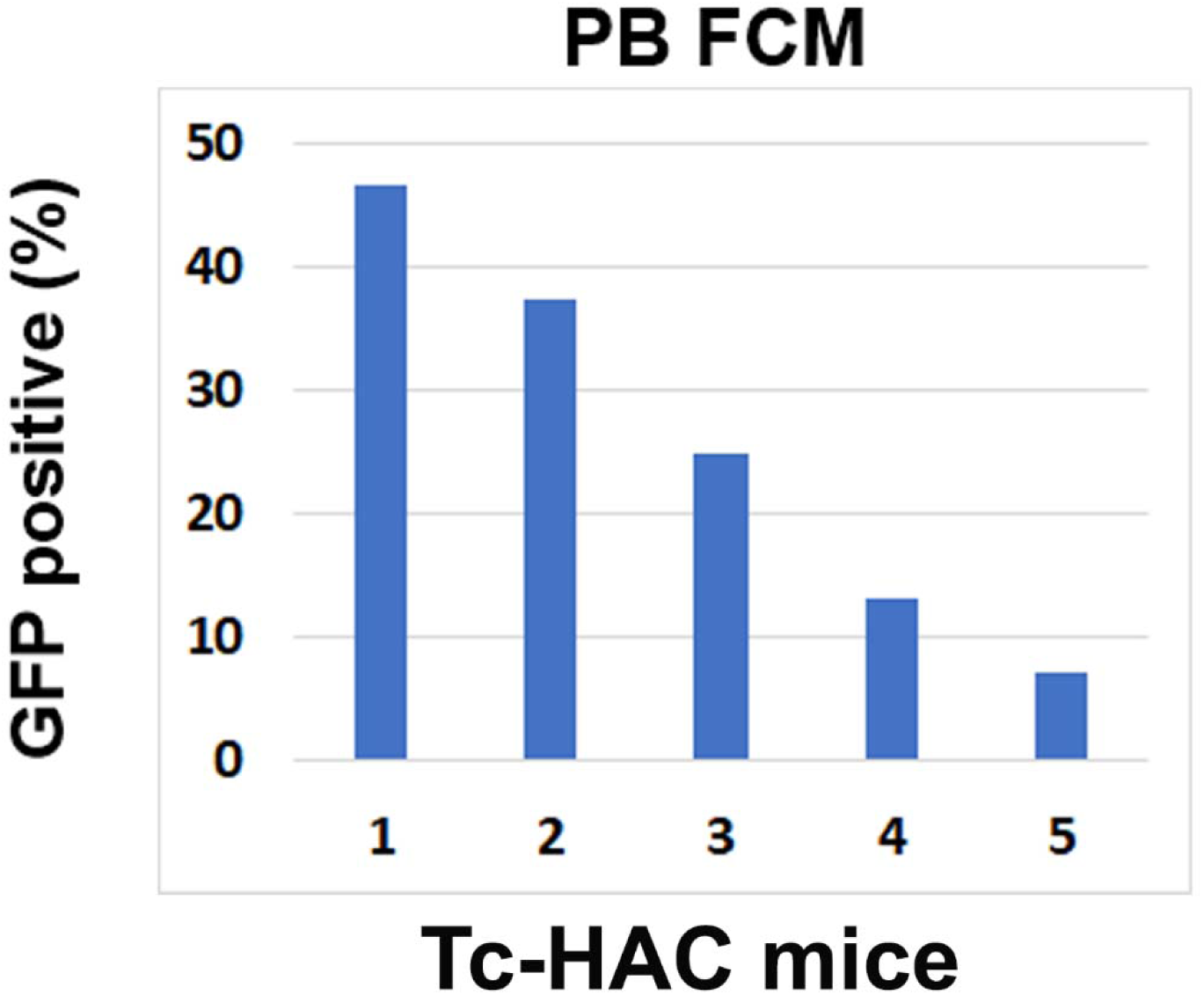
Mosaicism analysis of peripheral blood (PB) cells from Tc-HAC mice (n=5) using FCM.

