## Supplementary table 1,2,5,7,8 for "The first transchromosomic rat model with human chromosome 21 shows robust Down syndrome features"

### Slide 1
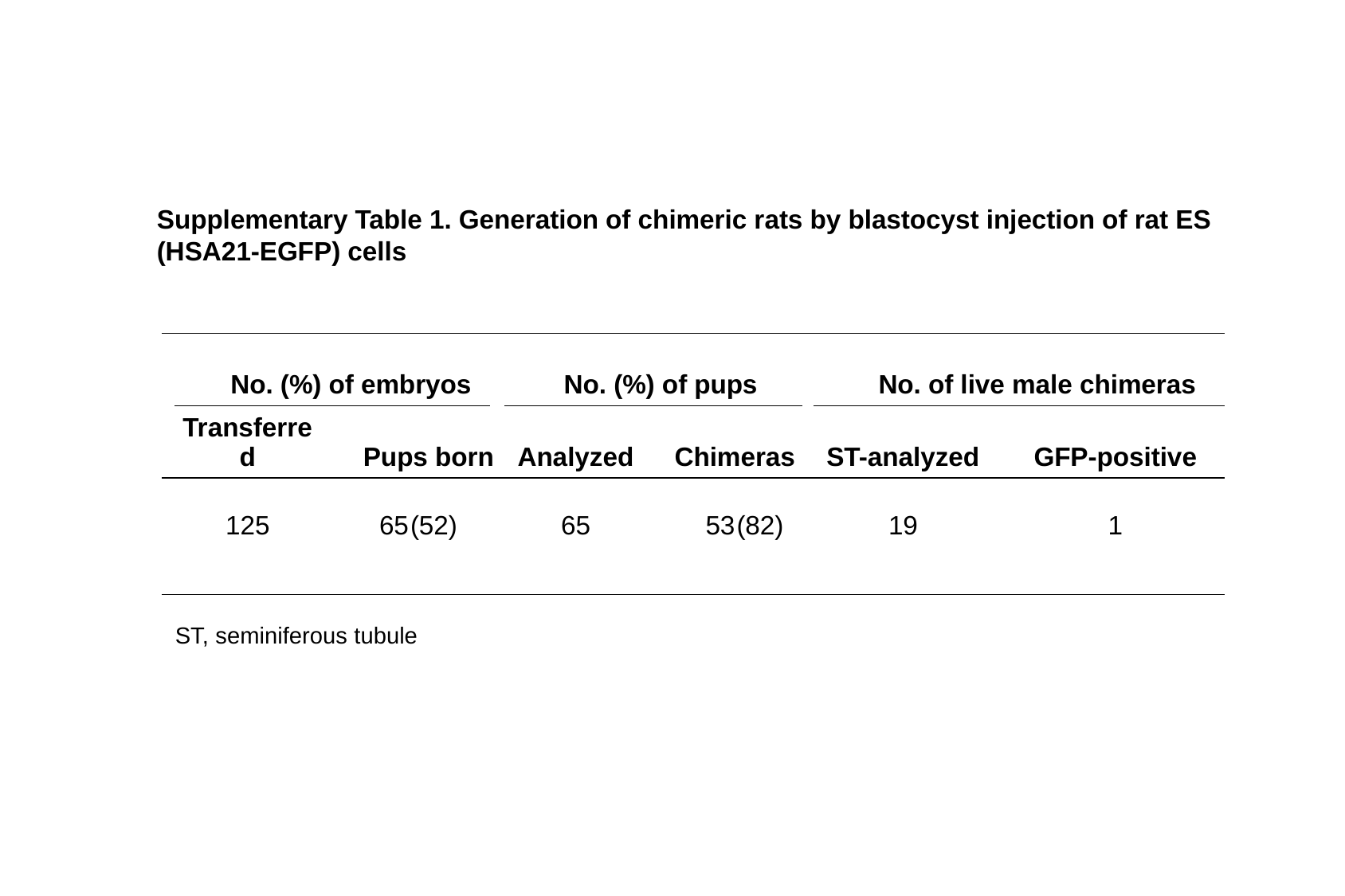

Supplementary Table 1. Generation of chimeric rats by blastocyst injection of rat ES (HSA21-EGFP) cells
| | | | | | | | | | | | | | | | | | | | | | |
| --- | --- | --- | --- | --- | --- | --- | --- | --- | --- | --- | --- | --- | --- | --- | --- | --- | --- | --- | --- | --- | --- |
| | No. (%) of embryos | | | | | | | | No. (%) of pups | | | | | | | | No. of live male chimeras | | | | |
| | Transferred | | | | Pups born | | | | Analyzed | | | Chimeras | | | | | ST-analyzed | | | | GFP-positive |
| | 125 | | | | 65 | (52) | | | 65 | | | 53 | | (82) | | | 19 | | | | 1 |
ST, seminiferous tubule

### Slide 2
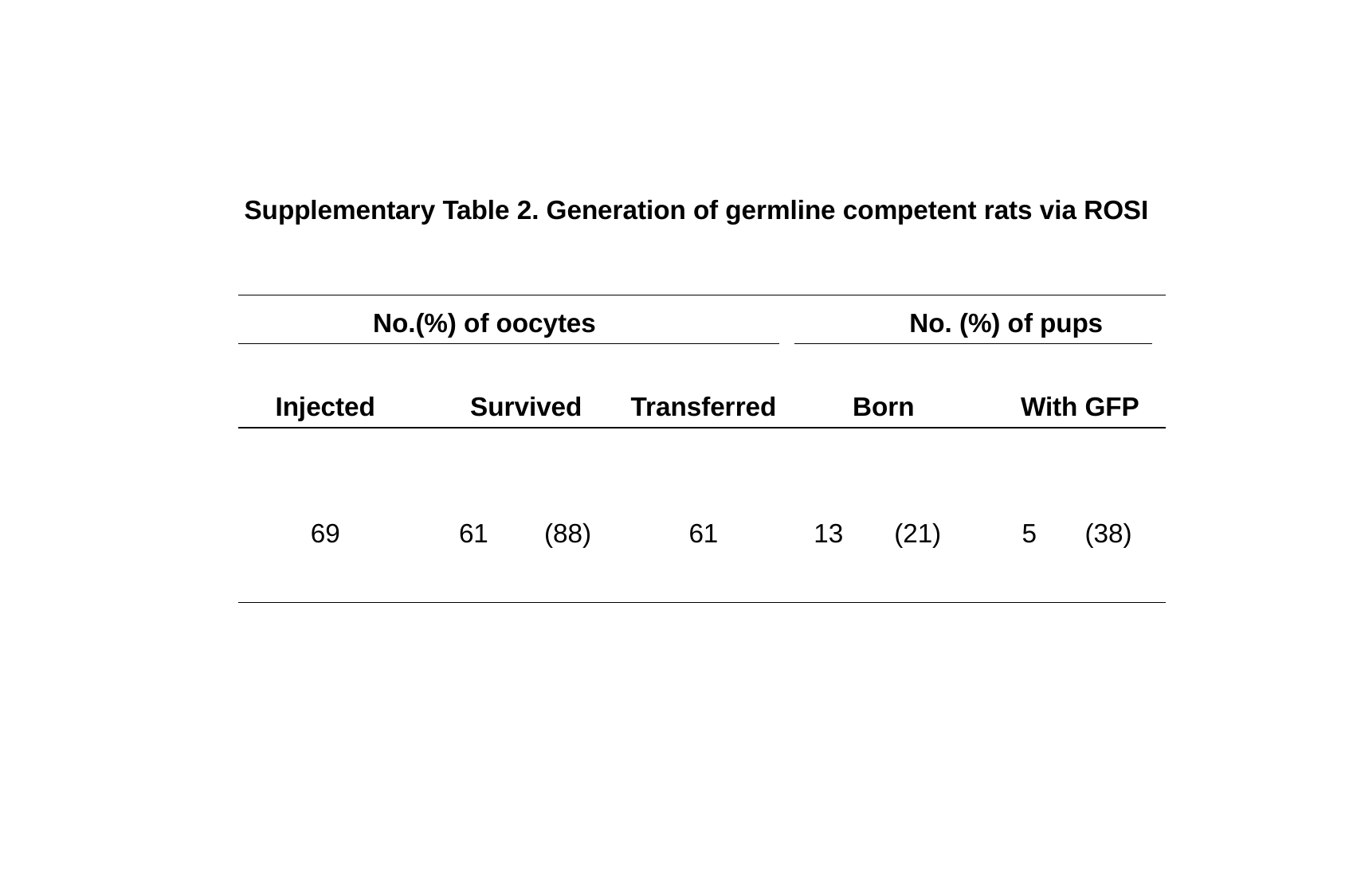

Supplementary Table 2. Generation of germline competent rats via ROSI
| | | | | | | | | | | | | | |
| --- | --- | --- | --- | --- | --- | --- | --- | --- | --- | --- | --- | --- | --- |
| No.(%) of oocytes | | | | | | | | No. (%) of pups | | | | | |
| Injected | | Survived | | | | Transferred | | Born | | | | With GFP | |
| 69 | | 61 | | (88) | | 61 | | 13 | (21) | | | 5 | (38) |

### Slide 3
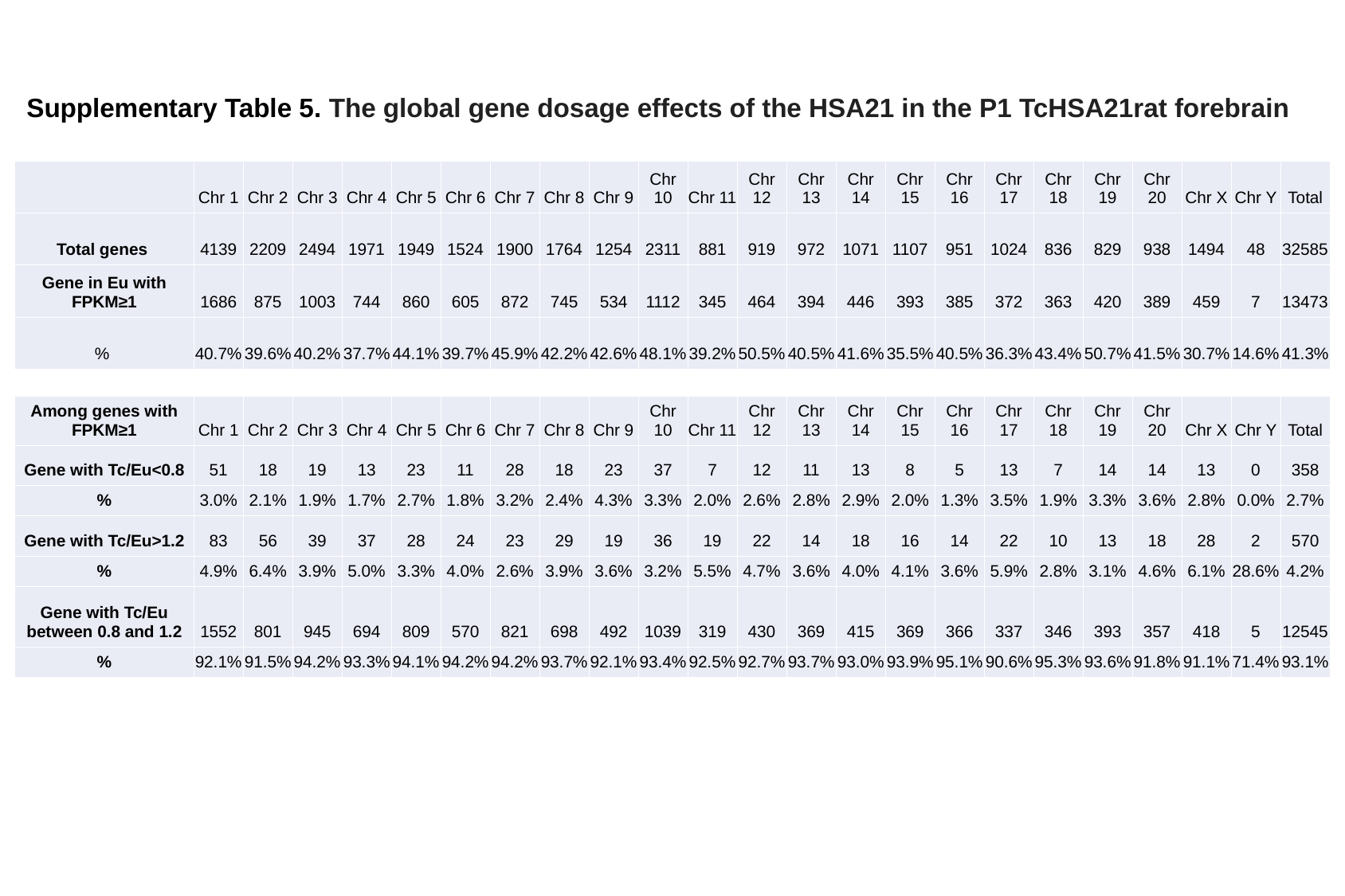

#### Supplementary Table 5. The global gene dosage effects of the HSA21 in the P1 TcHSA21rat forebrain
| | Chr 1 | Chr 2 | Chr 3 | Chr 4 | Chr 5 | Chr 6 | Chr 7 | Chr 8 | Chr 9 | Chr 10 | Chr 11 | Chr 12 | Chr 13 | Chr 14 | Chr 15 | Chr 16 | Chr 17 | Chr 18 | Chr 19 | Chr 20 | Chr X | Chr Y | Total |
| --- | --- | --- | --- | --- | --- | --- | --- | --- | --- | --- | --- | --- | --- | --- | --- | --- | --- | --- | --- | --- | --- | --- | --- |
| Total genes | 4139 | 2209 | 2494 | 1971 | 1949 | 1524 | 1900 | 1764 | 1254 | 2311 | 881 | 919 | 972 | 1071 | 1107 | 951 | 1024 | 836 | 829 | 938 | 1494 | 48 | 32585 |
| Gene in Eu with FPKM≥1 | 1686 | 875 | 1003 | 744 | 860 | 605 | 872 | 745 | 534 | 1112 | 345 | 464 | 394 | 446 | 393 | 385 | 372 | 363 | 420 | 389 | 459 | 7 | 13473 |
| % | 40.7% | 39.6% | 40.2% | 37.7% | 44.1% | 39.7% | 45.9% | 42.2% | 42.6% | 48.1% | 39.2% | 50.5% | 40.5% | 41.6% | 35.5% | 40.5% | 36.3% | 43.4% | 50.7% | 41.5% | 30.7% | 14.6% | 41.3% |
| Among genes with FPKM≥1 | Chr 1 | Chr 2 | Chr 3 | Chr 4 | Chr 5 | Chr 6 | Chr 7 | Chr 8 | Chr 9 | Chr 10 | Chr 11 | Chr 12 | Chr 13 | Chr 14 | Chr 15 | Chr 16 | Chr 17 | Chr 18 | Chr 19 | Chr 20 | Chr X | Chr Y | Total |
| --- | --- | --- | --- | --- | --- | --- | --- | --- | --- | --- | --- | --- | --- | --- | --- | --- | --- | --- | --- | --- | --- | --- | --- |
| Gene with Tc/Eu<0.8 | 51 | 18 | 19 | 13 | 23 | 11 | 28 | 18 | 23 | 37 | 7 | 12 | 11 | 13 | 8 | 5 | 13 | 7 | 14 | 14 | 13 | 0 | 358 |
| % | 3.0% | 2.1% | 1.9% | 1.7% | 2.7% | 1.8% | 3.2% | 2.4% | 4.3% | 3.3% | 2.0% | 2.6% | 2.8% | 2.9% | 2.0% | 1.3% | 3.5% | 1.9% | 3.3% | 3.6% | 2.8% | 0.0% | 2.7% |
| Gene with Tc/Eu>1.2 | 83 | 56 | 39 | 37 | 28 | 24 | 23 | 29 | 19 | 36 | 19 | 22 | 14 | 18 | 16 | 14 | 22 | 10 | 13 | 18 | 28 | 2 | 570 |
| % | 4.9% | 6.4% | 3.9% | 5.0% | 3.3% | 4.0% | 2.6% | 3.9% | 3.6% | 3.2% | 5.5% | 4.7% | 3.6% | 4.0% | 4.1% | 3.6% | 5.9% | 2.8% | 3.1% | 4.6% | 6.1% | 28.6% | 4.2% |
| Gene with Tc/Eu between 0.8 and 1.2 | 1552 | 801 | 945 | 694 | 809 | 570 | 821 | 698 | 492 | 1039 | 319 | 430 | 369 | 415 | 369 | 366 | 337 | 346 | 393 | 357 | 418 | 5 | 12545 |
| % | 92.1% | 91.5% | 94.2% | 93.3% | 94.1% | 94.2% | 94.2% | 93.7% | 92.1% | 93.4% | 92.5% | 92.7% | 93.7% | 93.0% | 93.9% | 95.1% | 90.6% | 95.3% | 93.6% | 91.8% | 91.1% | 71.4% | 93.1% |

### Slide 4
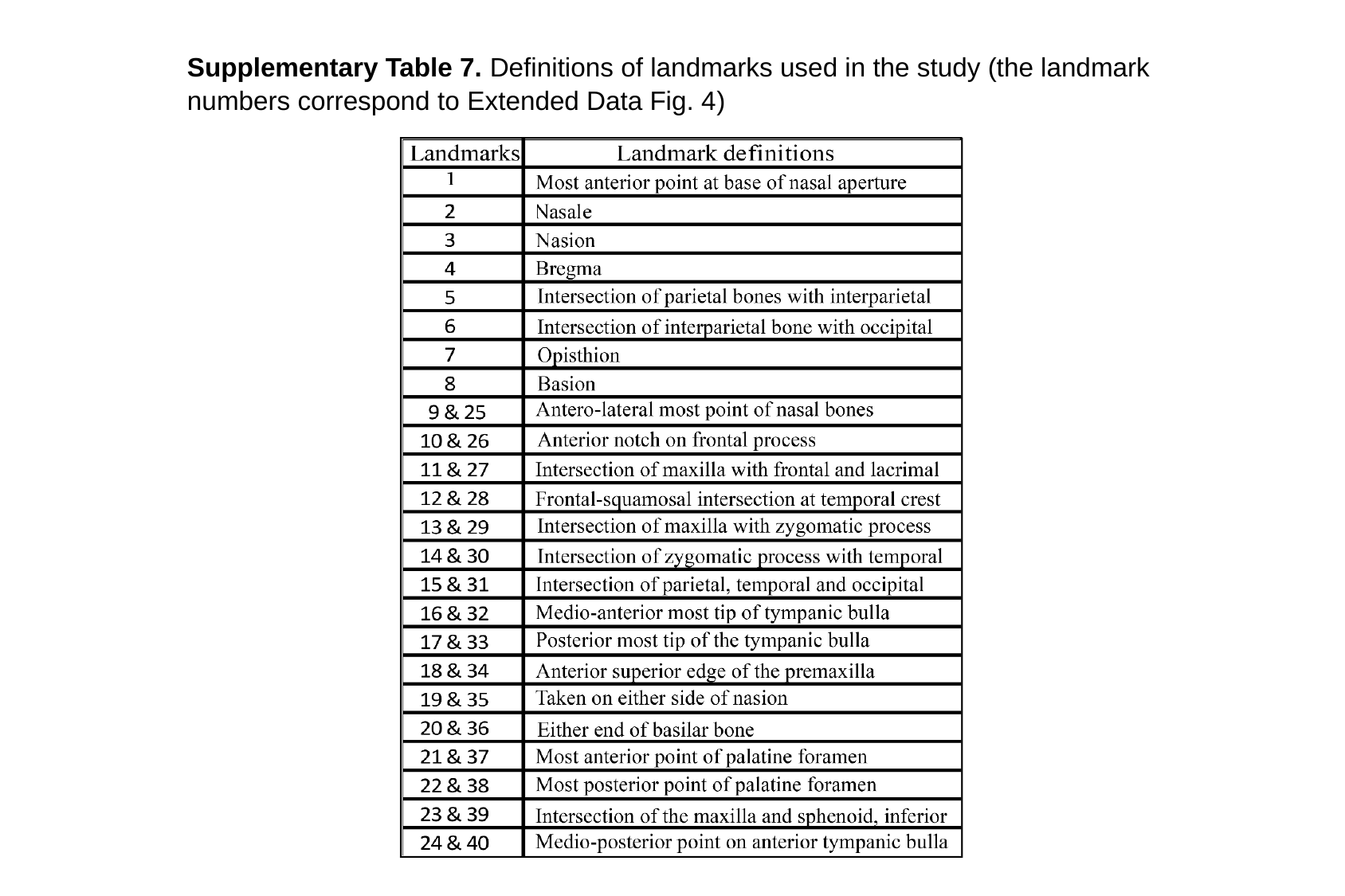

Supplementary Table 7. Definitions of landmarks used in the study (the landmark numbers correspond to Extended Data Fig. 4)

### Slide 5
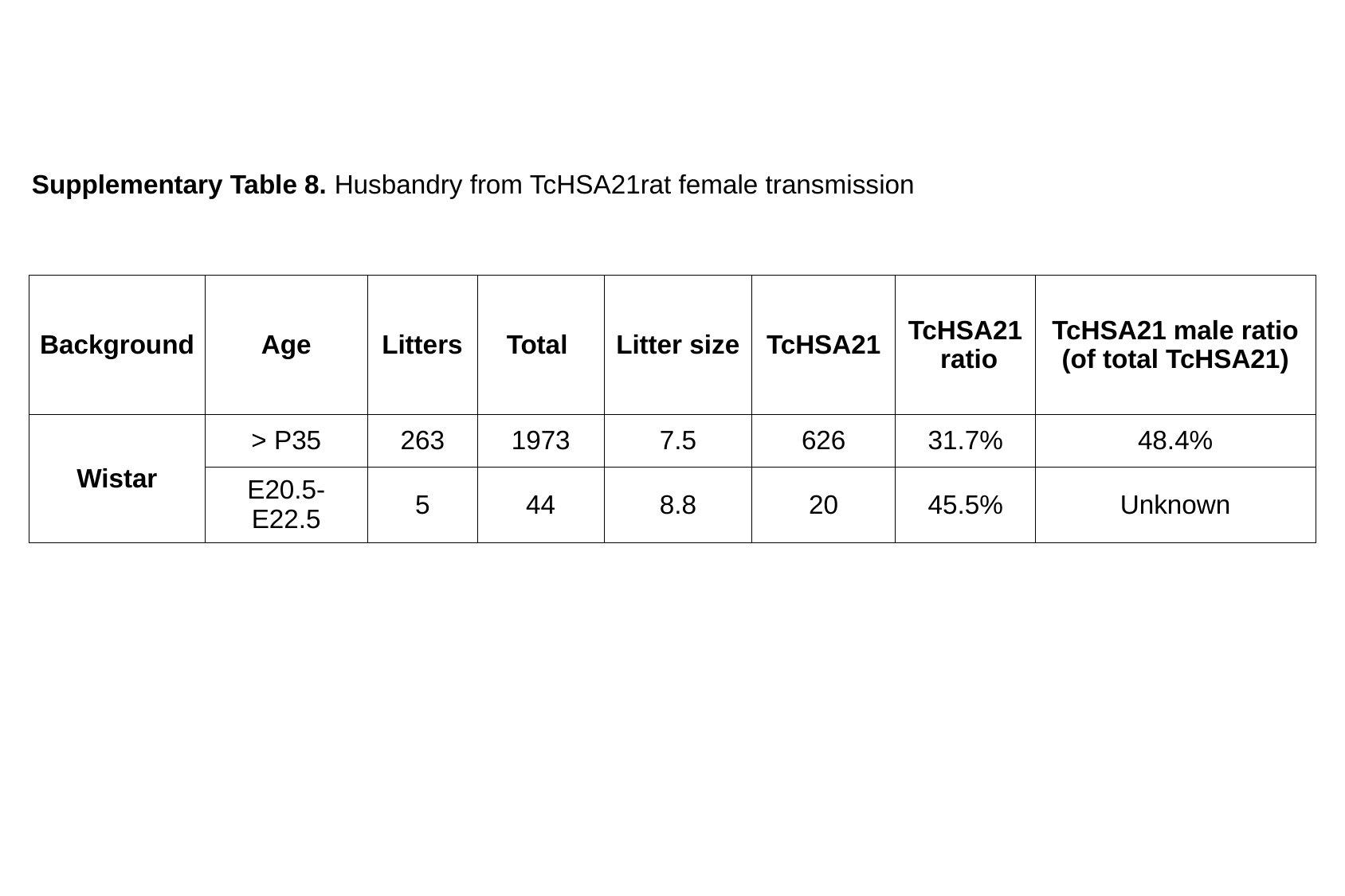

Supplementary Table 8. Husbandry from TcHSA21rat female transmission
| Background | Age | Litters | Total | Litter size | TcHSA21 | TcHSA21 ratio | TcHSA21 male ratio (of total TcHSA21) |
| --- | --- | --- | --- | --- | --- | --- | --- |
| Wistar | > P35 | 263 | 1973 | 7.5 | 626 | 31.7% | 48.4% |
| | E20.5-E22.5 | 5 | 44 | 8.8 | 20 | 45.5% | Unknown |
